# Sequencing red fox Y chromosome fragments to develop phylogenetically informative SNP markers and glimpse male-specific trans-Pacific phylogeography

**DOI:** 10.1101/552323

**Authors:** BN Sacks, ZL Lounsberry, HM Rando, K Kluepfel, S Fain, SK Brown, AV Kukekova

## Abstract

The red fox (*Vulpes vulpes*) has a wide global distribution with many ecotypes, and has been bred in captivity for various traits, making it a useful evolutionary model system. The Y chromosome represents one of the most informative markers of phylogeography, yet it has not been well-studied in the red fox due to a lack of the necessary genomic resources. We used a target capture approach to sequence a portion of the red fox Y chromosome in a geographically diverse red fox sample, along with other canid species, to develop single nucleotide polymorphism (SNP) markers, 13 of which we validated for use in subsequent studies. Phylogenetic analyses of the Y chromosome sequences, including calibration to outgroups, confirmed previous estimates of the timing of two intercontinental exchanges of red foxes, the initial colonization of North America from Eurasia approximately half a million years ago and a subsequent continental exchange before the last Pleistocene glaciation (∼100,000 years ago). However, in contrast to mtDNA, which showed unidirectional transfer from Eurasia to North America prior to the last glaciation, the Y chromosome appears to have been transferred from North America to Eurasia during this period. Additional sampling is needed to confirm this pattern and to further clarify red fox Y chromosome phylogeography.

---

The phylogeography of red fox (*Vulpes vulpes*) has been well-characterized in terms of mitochondrial and nuclear patterns of diversity over most of its global distribution (Aubry et al. 2009; Edwards et al. 2012; Kutschera et al. 2013; Statham et al. 2014, 2018; Goldsmith et al. 2016; Sacks et al. 2018). For example, the most divergent mitochondrial split dates to approximately half a million years ago and separates a clade that evolved in North America south of the ice sheets prior to the last glaciation (Nearctic clade) and one that spans Eurasia and Alaska/western Canada, reflecting a secondary continental exchange event ∼100 thousand years ago (ky; Aubry et al. 2009; Statham et al. 2014). However, little is known about red fox Y chromosome diversity. Except for microsatellites, no markers exist for such investigations in red foxes (Statham et al. 2014; Rando et al. 2017). Although microsatellites are useful for population-genetic questions about contemporary gene flow and recent historical demography, more conserved mutations are needed for deeper reconstruction of Y chromosome phylogenies.

Sequencing the Y chromosome is challenging, even in model species, due to an evolutionarily plastic history that has resulted in high frequency of repetitive DNA, paralogs of autosomal and X chromosomal regions, and palindromes (Charlesworth and Charlesworth 2000; Skaletsky et al. 2003; Tomaszkiewicz et al. 2016). The challenge is even more pronounced in non-model organisms due to a lack of reference genomes. Notwithstanding these obstacles, it is feasible to enrich libraries for DNA resembling Y chromosome sequence from other mammals, sequence these enriched libraries, and use read depth as a criterion for distinguishing unique loci from repetitive motifs (Tomaszkiewicz et al. 2016; Rangavittal et al. 2018). As a final confirmation, primers can be designed to genotype the presumptive unique loci and to test them in male and female individuals (Natanaelsson et al. 2006). Those that turn out to be male-specific and provide no more than one allele per individual can be inferred to be male-specific Y chromosome markers. Because the entire male-specific length of the Y chromosome is linked, these markers can be employed in tandem, regardless of their relative locations on the chromosome, as multi-SNP haplotypes to reconstruct phylogenetic topologies.

In this study, we used information from the dog genome to enrich genomic fox libraries for Y chromosome DNA and sequenced the libraries in a diverse geographic sample of red fox to develop Y chromosome SNP markers and provide a preliminary look at male-specific phylogeography. Although dogs and red foxes are relatively close relatives, they have distinct karyotypes, and sex chromosomes in particular are evolutionarily unstable (Charlesworth and Charlesworth 2000). Therefore, we emphasized use of Y chromosome regions that were conserved across multiple orders of mammal (Natanaelsson et al. 2006) and validated SNP markers on male and female red foxes. To obtain a red fox Y chromosome tree and divergence time estimates for major Y chromosome clades, we conducted a phylogenetic analysis rooting and calibrating branch lengths among red fox Y chromosome sequences, those of the most basal extant canids (*Urocyon* spp.), and of coyotes (*Canis latrans*).

## Materials and Methods

### Samples

For high-throughput sequencing, we used 18 male red fox samples (Fig. 1), along with 2 gray foxes (*Urocyon cinereoargenteus*), an island fox (*U. littoralis*), and 6 coyotes (*Canis latrans*) as outgroups (Supplementary Table S1). These samples were used for SNP discovery and phylogenetic analyses. We used 9 of the same red fox males, one additional red fox male from Alaska, and 19 red fox females for red fox SNP marker validation.

**Figure 1.**
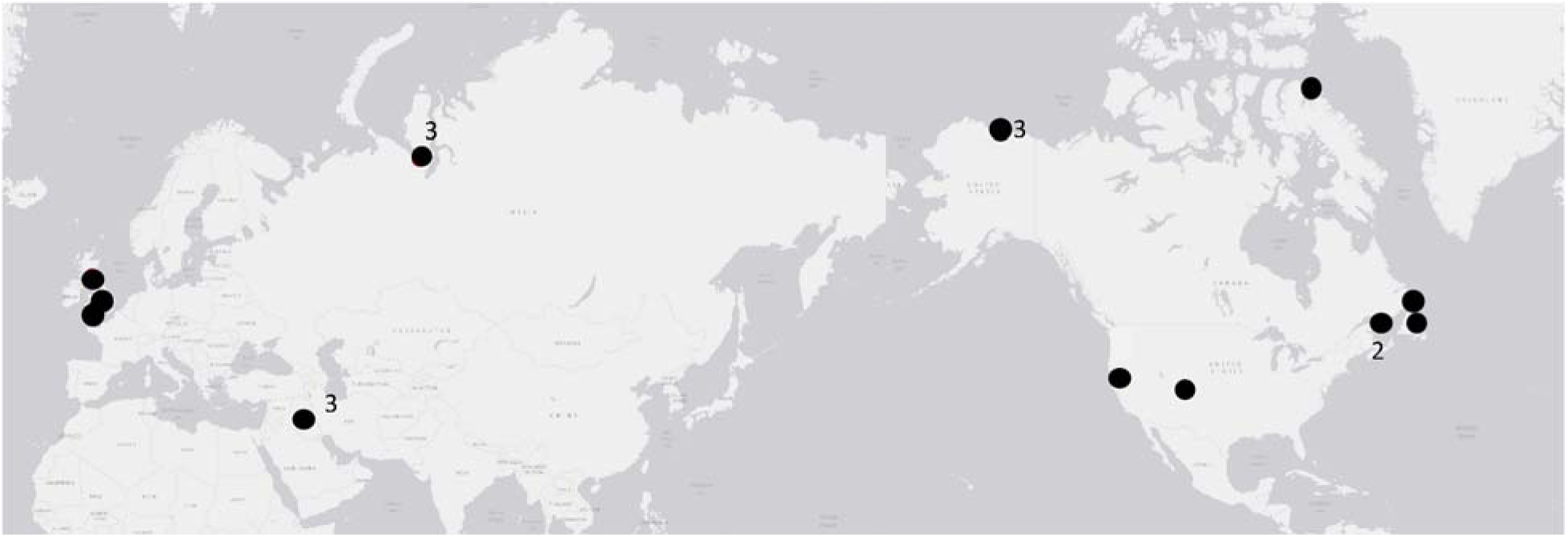
Locations of male red fox samples used in this study (n = 19). One sample from Alaska was genotyped for selected SNPs but not sequenced. Numbers refer to the sample size indicated by a marker. Markers with no numbers are a single sample.

### Capture enrichment and sequencing

We constructed individually barcoded whole genome shotgun libraries using the NEBNext Ultra DNA Library Prep Kit for Illumina (New England Biolabs, Ipswich, MA) and 6-bp custom indexed primers. We then pooled libraries (10 ng each) in batches of 47 samples for hybridization to capture baits. Although the entire dog Y chromosome assembly is now available (Li et al. 2013), we began this study just prior to its publication and, therefore, relied on a smaller subset of canine Y chromosome fragments totaling 39,246 bp. (However, we did map markers to the Li et al. 2013; GenBank Accession No. KP081776.1.) We designed baits that were complementary to previously published and validated dog Y-chromosome DNA sequences (Supplementary Information 1) that had been prescreened to avoid paralogy to X chromosome sequences from canids and other mammals (Hellborg and Ellegren 2003; Natanaelsson et al. 2006; Ding et al. 2012; Tsubouchi et al. 2012). We also included capture baits for exons from the Y chromosome amelogenin (AMELY) and zinc finger (ZFY) genes, which do have high paralogy with the X chromosome, but could have unique regions as well. Baits were each 80 bp in length and overlapped adjacent baits by 40 bps (i.e., tiling density = 2x), as appropriate for capture between closely related species. The RNA baits were synthesized and biotinylated by MyBaits (MYcroarray, Ann Arbor, MI). We conducted the hybridization with blockers for 36 hours at 65°C according to manufacturer’s protocols, using the MyBaits Target Enrichment Kit (Arbor Biosciences, Ann Arbor, MI, USA) We bound hybridization products to streptavidin-coated magnetic beads, washed away unbound library, isolated bound (i.e., enriched) library from baits and beads, and PCR-amplified the enriched library for 14 cycles. We then sequenced the pooled enriched library on an Illumina MiSeq lane SR300 at the UC Davis Genome Center. Additionally, we included HiSeq2500 reads (PE 150) from 2 male red fox whole genome shotgun libraries sequenced for a different project.

### Bioinformatic processing

We trimmed adapter sequences and filtered reads for quality, discarding reads with >50% of bases with quality scores <2, and trimmed low-quality 3’ end bases using Next Generation Sequencing Short Read Trimmer (ngsShoRT; Chen *et al.* 2014). We aligned reads to the original bait sequences using bwa-mem (Li and Durbin 2009) and processed alignments with Samtools (Li et al. 2009). We then used freeBayes v0.9.21 (Garrison and Marth 2012) to call variants against the original bait sequence references, allowing “heterozygotes” to flag paralogs downstream, and exported all positions for each individual (variable and not) in variable call format (vcf). Because we had no independent knowledge of X-degenerate (uniquely mapping) red fox or *Urocyon* Y chromosome sequence, we used coverage depth as a basis for selecting positions empirically. The distribution of coverage depths within a species contained multiple modes, which we presumed corresponded on the low end to erroneous sites (e.g., exogenous contamination), followed by uniquely aligning sites, sites aligning to both sex chromosomes, and, lastly, a long tail corresponding to variable frequency repetitive motifs. Therefore, after visualizing the distributions for red foxes, we selected a coverage-depth range corresponding to the second mode and either tail (see Results). We used these sites to obtain intraspecific SNPs.

### Phylogenetic analysis

We further filtered the sites inferred above to be uniquely aligning to the red fox Y chromosome based on the same criteria described above. We also removed any interspecific indels to obtain Y chromosome sequence orthologous across all taxa for calibrating substitution rates against independently determined node ages (Wayne et al. 1997; Lindbladh-Toh et al. 2005; Perini et al. 2009). We used Mega 6 (Tamura et al. 2013) to construct a maximum likelihood tree with 500 bootstrap replicates based on the Tamura 3 parameter mutation model (T92; Tamura 1992), which was supported over 23 other models by the lowest Bayesian Information Criterion. To estimate divergence time between red fox clades, we computed pairwise T92 distances and standard errors between red fox clades and calibrated these to pairwise distances to coyote and gray fox, previously estimated to be 8.8 MY divergent from red foxes (Perini et al. 2009).

### SNP assay

We used Assay Design Suite 2.0 (https://mysequenom.com/Tools) to design primers for multiplexing on the MassARRAY iPLEX platform (Agena Biosciences, Inc., San Diego, CA). To validate these SNPs as uniquely representing the non-recombining portion of the Y chromosome, we tested them in both male and female foxes and retained only those yielding male-specific genotypes.

### Intraspecific red fox network

After combining sequencing and genotyping calls, a small number of remaining missing calls were imputed parsimoniously with the allele found in otherwise identical complete sequences. We constructed statistical parsimony network using the median joining method in Networks (v 5.0; Fluxus Technology Ltd, England; Bandelt et al. 1999; Forster et al. 2000) and estimated the age of clades in terms of the average (and standard error) numbers of mutations separating derived from inferred ancestral nodes (Forster et al. 2000; Saillard et al. 2000).

## Results

Excluding one red fox sample that failed sequencing, we obtained 6,486,865 raw 300-bp reads from a MiSeq lane of the 23 target-enriched samples of which 2,393,840 (36.8%) reads aligned back to the original bait sequences (i.e., 41 targeted fragments of the dog Y chromosome). This total included 1,342,241 of 3,304,125 reads (40.6%) from 14 red foxes, 738,554 of 2,296,795 reads (32.2%) from 6 coyotes, and 313,045 of 898,307 reads (34.8%) from 3 *Urocyon* foxes. We also obtained 86,687,990 bp of HiSeq sequence for 2 additional red foxes, of which 1,835,416 bp (2.1%) aligned to the bait sequences. The ratio of targeted sequence obtained from the capture enrichment vs whole genome shotgun approaches for red foxes (i.e., 40.6% vs 2.1%) indicate that the enrichment process increased yield approximately 20-fold over random sequencing.

In total, we recovered 37,080 sites, of which 31,624 (85.3%) had coverage depths in the range we presumed corresponded to X-degenerate sites on the Y chromosome (10–160×) and no heterozygous sites in red foxes (Fig. 2; vcf file deposited in Dryad). For phylogenetic analyses, we retained 20,931 of these sites after removing interspecific indels and confirming that read depths in the other species were also consistent with uniquely aligning sites. These 20,931 sites had an average of 85× depth in red foxes (5.3× per individual), 85× in coyotes (14.3× per individual), and 30× in *Urocyon* spp. (10× per individual).

**Figure 2.**
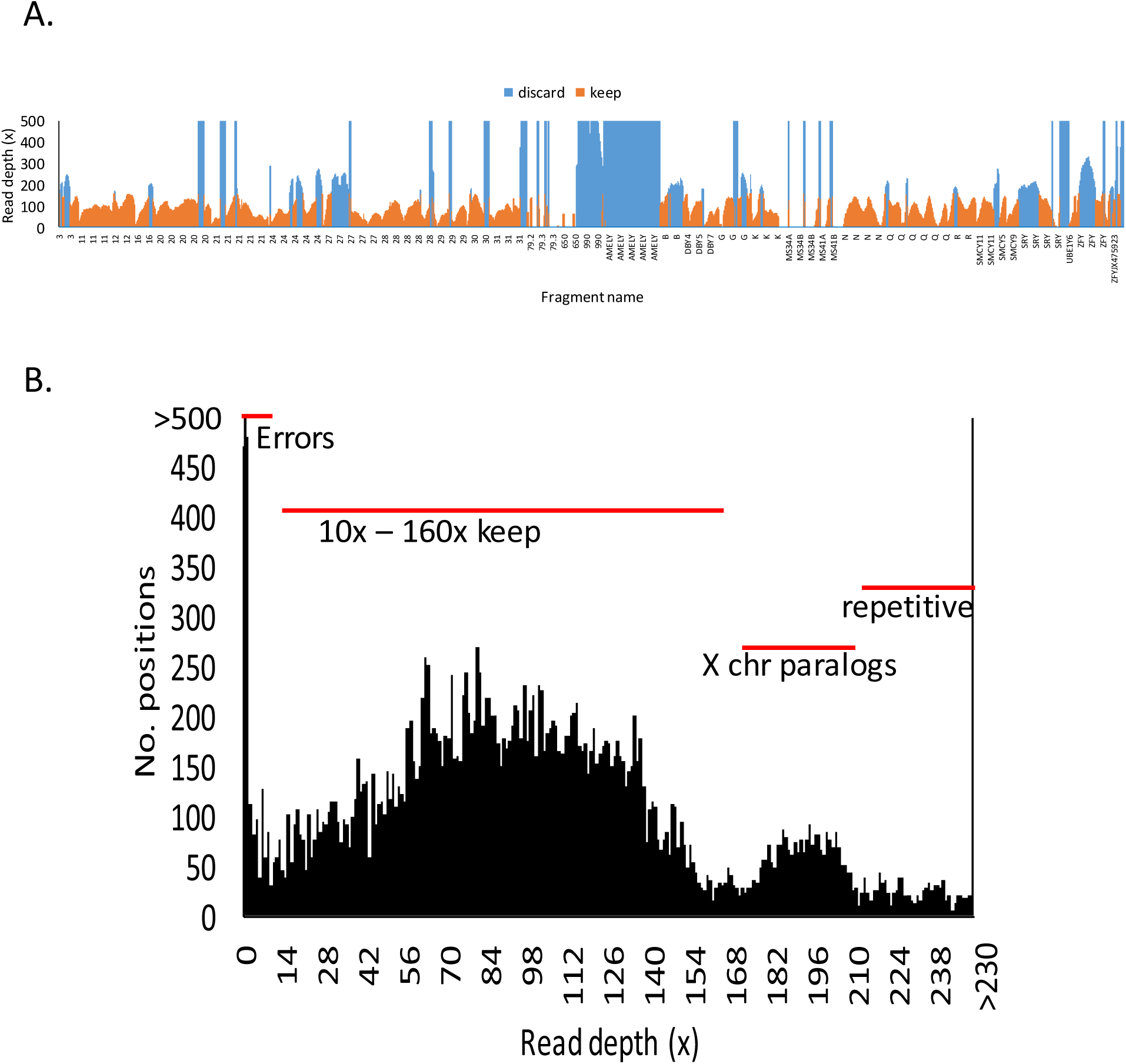
Coverage depths from sequencing reads of 19 male red foxes mapped to 37,080 sites across 41 orthologous dog Y chromosome fragments shown (A) with read depths arrayed across ordered sites on the reference fragments, and (B) as a frequency distribution, illustrating a systematic pattern of coverage presumably corresponding to uniquely mapping Y chromosome sites (10-160x), sites likely to reflect 2 paralogs (e.g., including one on the X chromosome, 161-210x), and sites with >2 paralogs (>210x). Only sites with depths ranging 10-160x were used in analyses. Sites with depths <10x were presumed to reflect errors and those >160x were presumed to reflect paralogs, not necessarily all on the Y chromosome.

In the 20,931 site dataset (all taxa), we observed 460 substitutions, including 54 intraspecific (or intrageneric in the case of *Urocyon*) polymorphisms and 404 interspecific substitutions (254 Ti, 150 Tv). Two of the 54 intraspecific polymorphisms occurred in both red foxes and *Urocyon*, altogether yielding 22 polymorphisms (14 Ti, 8 Tv) in 17 red foxes, 13 polymorphisms (9 Ti, 4 Tv) among the 3 *Urocyon* foxes, and 23 polymorphisms (14 Ti, 9 Tv) among the 6 coyotes. A maximum likelihood tree indicated two well-supported, reciprocally monophyletic red fox clades (Fig. 3). Based on the Tamura 3 parameter model distances (and SE) calibrated to *Urocyon* and coyote divergence times (8.8 MY, Perini et al. 2009), the two red fox Y chromosome clades diverged approximately 470,000 (402,000–563,000) years ago, which was consistent with mitochondrial and autosomal estimates (Statham et al. 2014).

**Figure 3.**
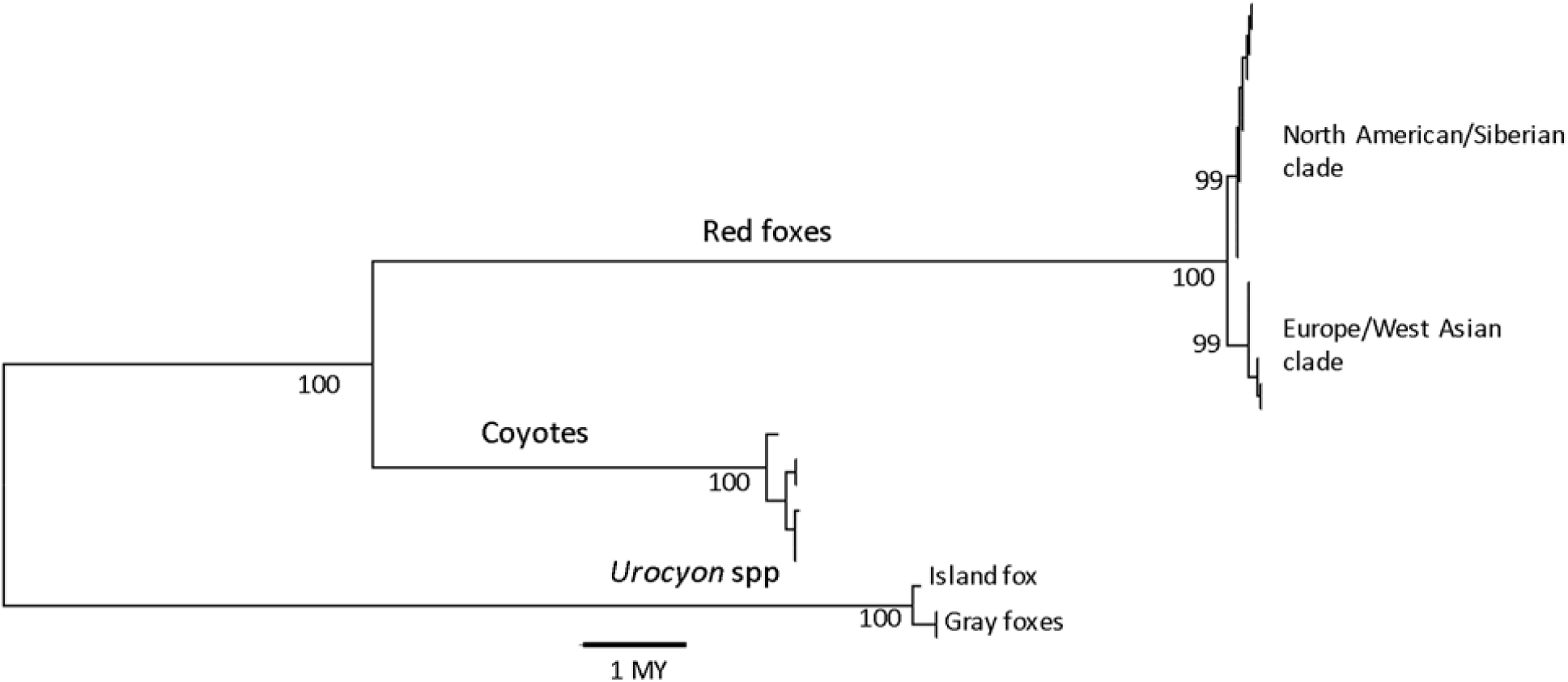
Red fox Y chromosome phylogeny displayed as Maximum Likelihood tree inferred from 20,931 bp of sequence in 17 red foxes, 2 gray foxes, an island fox, and 6 coyotes with node support inferred from 500 bootstrapped trees. Genetic distance was computed according to the Tamura 3-parameter model (Tamura 1992) and calibrated to divergence time in millions of years (MY).

**Figure 4.**
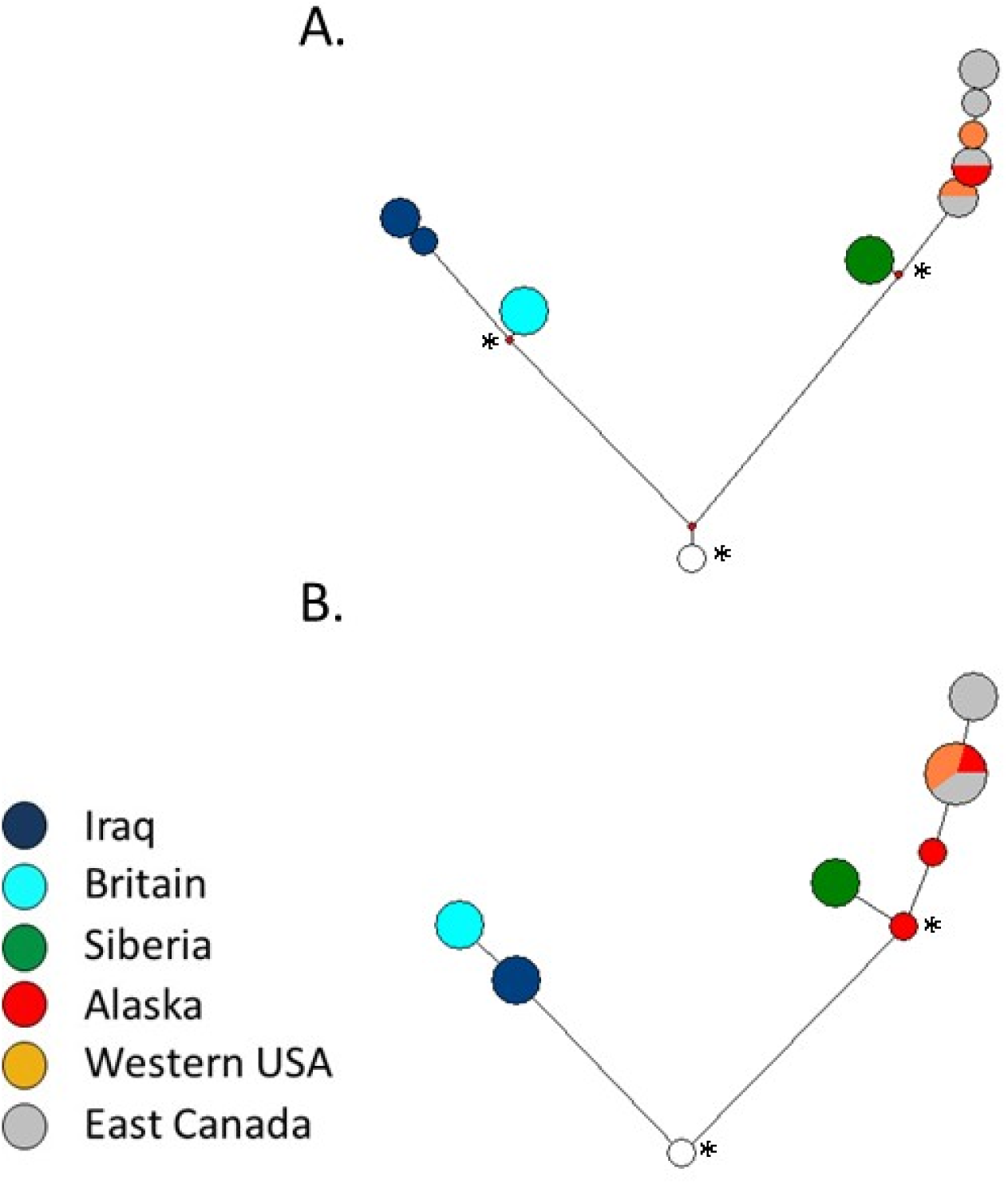
Median joining networks utilizing (A) 31 Y chromosome SNPs in 17 red foxes or (B) a subset of 13 assayed Y chromosome SNPs in 19 red foxes (includes 2 additional Alaskan red foxes); the ancestral node (or root, white circle) in each network corresponded to the haplotype common to all coyotes, gray foxes, and the island fox. Red dots in (A) represent inferred nodes. (A,B) Nodes (observed or inferred) used for aging haplogroups through rho statistics are indicated with an asterisk. Node sizes are proportional to number of foxes and branch lengths are proportional to the number of mutations.

For intraspecific analyses of red foxes, we retained all of the 31,624 presumptive Y chromosome sites. Among these sites, we observed 31 SNPs within red foxes (including the 22 intraspecific sites used above, along with 9 additional sites); we designed SNP assays for 14 of them (Table 1; Supplementary Tables S2, S3). We tested SNP assays on 9 of the originally sequenced male red foxes and 19 female red foxes and also genotyped the Alaskan male that failed sequencing and one other Alaskan male for which no sequencing was attempted. All genotypes for the 9 males previously sequenced matched those based on the original sequences. All except one of the loci failed to amplify consistently in females. Locus 20_357AT amplified in 14 of 19 female samples, indicating that the primers we designed were not sufficiently specific to the Y chromosome target. Otherwise, the false-positive rate among the other 13 loci was 3.5% in females, with no single locus or individual yielding notably more than any other. Positions of the 13 validated SNPs were determined based on the Y chromosome dog assembly (Supplementary Table S3).

**Table 1.**
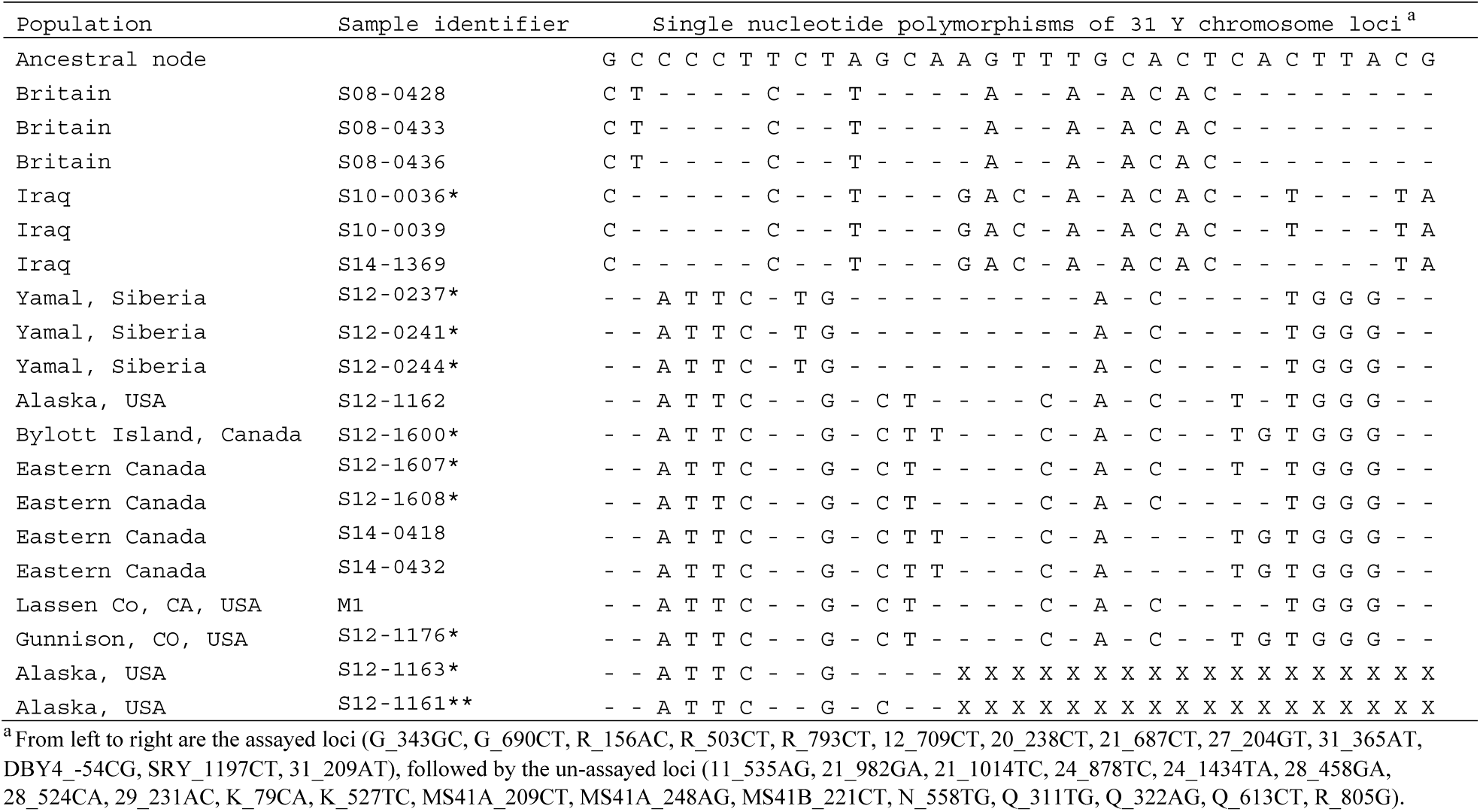
Y chromosome haplotypes of 19 male red foxes and an ancestral node inferred from orthologous positions in *Canis* and *Urocyon*, which are composed of 31 variable sites discovered 31,624 bp of Y chromosome sequence. We also designed a genotyping assay for 13 of the loci (first 13 from left to right). Eighteen male red foxes were sequenced, including 9 foxes that were also genotyped (*), and 1 fox was genotyped but not sequenced (**). One locus (20_349AT) that amplified in 14 of 19 females was excluded from this table. None of the other 13 genotyped loci consistently amplified in females, although we observed a low (3.5%) false-positive rate. Genotype and sequencing calls matched 100% of the time. Two foxes that were genotyped but not sequenced have missing data (X) across all non-assayed sites.

We constructed two median joining networks: one using all 31 variable sites in the 17 successfully sequenced red foxes (Fig. 3A), and one using the subset of 13 assayed and validated sites that additionally included 2 Alaskan male red foxes genotyped using the assay (Fig. 3B; Table 1). The former network provided more resolution of haplotypes, particularly within continents, whereas the latter network included two additional Alaskan foxes with haplotypes that were basal within their clade. For the network based on all 31 sites, the rho estimates (i.e., average numbers of mutations since divergence from the most recent common ancestor) were 13.00 (SD = 2.55) for the entire network and 3.82 (SD = 1.48) for the North American/Siberian clade, indicating a ratio of 0.294. For the network based on 13 sites, the rho estimates were 4.80 (SD = 1.55) for the entire network and 1.77 (SD = 0.98) for the North American/Siberian clade, indicating a ratio of 0.369. Assuming a 470,000 year divergence between the two clades, these estimates imply an age of the most recent common ancestor of Siberian and North American red foxes ranging 77,000 to 270,000 years ago based on the 13-site network or 85,000 to 190,000 years ago based on the 31-site network, both of which are consistent with previous mtDNA and autosomal nuclear estimates (Statham et al. 2014).

## Discussion

The Y chromosome is replete with repetitive elements and regions paralogous to other chromosomes, posing several significant obstacles to sequencing even in species for which the Y chromosome genome has been assembled and provides a reference (Tomaszkiewicz et al. 2016). Our reliance on dog Y chromosome sequence, which likely shared only partial homology to red and gray foxes, complicated this task further. Nevertheless, our efforts to enrich fox libraries for Y chromosome reads and obtain sequence for non-repetitive fox Y chromosome fragments were successful.

Overall, we obtained approximately 75% of the targeted Y chromosome sequence as uniquely aligning in male red foxes. Although 1 of the 14 SNPs tested in females yielded regular amplification products, this may have been due to the non-specificity of the primers themselves rather than of the contiguous sequence reflected in the sequencing reads. The read depth at this SNP in the original sequencing reads was 132× among red foxes, which was well within the range of depths observed in most positions and substantially smaller than the range corresponding to a larger mode and presumed to correspond to loci with X and Y paralogs. Regardless, 13 of the loci for which we developed a genotyping assay resolved two major clades, as well as several haplotypes within each. We also discovered 18 additional polymorphisms for which SNP assays could be developed and validated in the future. Most of these 18 SNPs would further differentiate the already well-defined major clades (Table 1). More importantly, 3 intraspecific polymorphisms (i.e., 29_231AC, MS41A_209CT, MS41A_248AG) may be useful in further resolving North American haplotypes.

Given the nearly global distribution of red foxes and many translocations (Long 2003), these markers will be useful in complementing mitochondrial markers as a way of easily identifying continent of origin, for example, in the case of introductions (e.g., Kasprowicz et al. 2016). Moreover, combining these SNPs with Y-linked microsatellites (Rando et al. 2017) can elucidate historical demographic patterns on Holocene time scales (Forster et al. 2000: Sacks et al. 2013).

Although our study was not designed primarily as a phylogeography study, which would require many more samples, our findings nevertheless provided some insights about the red fox’s continental Y chromosome phylogeography in the context of previous models of continental exchange between Eurasian and North American red foxes. Most fundamentally, our Y chromosome phylogeny provided independent support for previous estimates of the timing of continental exchanges between red foxes, specifically, an initial colonization of North America by Eurasian red foxes approximately 500 ky and a secondary continental exchange around the beginning of the last glaciation ∼100 ky (Aubry et al. 2009; Statham et al. 2014).

Prior to this study, discordance observed between mitochondrial and nuclear genetic patterns resulted in unresolved hypotheses about the nature and extent of the secondary exchange (Statham et al. 2014; Sacks et al. 2018). In particular, nearly all of the mitochondrial haplotypes observed in modern red foxes from Alaska and much of northwest Canada traced to Eurasia around the time of the secondary exchange (e.g., Aubry et al. 2009; Statham et al. 2014; Goldsmith et al. 2016). In contrast, most of the nuclear genetic ancestry in this same region traced to the original colonization of North America 500 ky, with little genetic exchange in either direction corresponding to the secondary contact event (Statham et al. 2014; Sacks et al. 2018). These discordant patterns suggested two hypotheses: (1) a selective sweep on Eurasian mtDNA introduced into North American red foxes following the secondary exchange, or (2) a male-mediated expansion of southern North American red foxes after the recession of the two major North American ice sheets after the last glacial maximum (<20 ky), leading to partial replacement of a Beringian population formerly composed of Eurasian red foxes. These hypotheses differ in whether the pre-event (continental exchange, expansion) population in Beringia was composed of Eurasian or North American red foxes and in the timing of the event that led to the discordance.

In the present study, we found the major split in the Y chromosome phylogeny dated to 500 ky, presumably reflecting the same secondary intercontinental split seen in nuclear and mitochondrial data. However, one of the clades included all North American, as well as Siberian, samples. The estimated age of the most recent common ancestral haplotype of Siberian and North American haplotypes dated to approximately 100 ky, consistent with Y chromosome movement from North America to Eurasia during the intercontinental exchange. This observation is consistent with hypothesis 1 but not hypothesis 2, which implies North American patrilines did not occur in Beringia until after the second continental exchange ∼100 ky. Thus, the exclusivity of Eurasian mitochondrial haplotypes combined with the general rarity of Eurasian nuclear ancestry in modern-day Alaskan red foxes most likely traces to a selective sweep ∼100 ky on the mitochondrial genome.

Clearly our sample size was too small to address the extent to which North American Y chromosomes contributed to East Asian red fox diversity, or whether some reciprocal exchange could have occurred. These questions can be investigated further in the future by genotyping larger samples of foxes with the SNPs developed in this study, and in combination with Y chromosome microsatellite markers (Statham et al. 2014; Rando et al. 2017).

## Supplementary Material

Supplementary data are available at Journal of Heredity online.

## Acknowledgments

We thank H Liu for assistance in sample preparation. We thank B. Popper, B. Chomel, C. Soulsberry, C. Miller, D. Berteaux, D. Ehrich, E. Tiller, J. Velasquez, J. Amaral, J. Perrine, J. Lindeman, M. Reid, P. Magee, P. Cross, P. Cervantez, S. Lariviere, T. Brinkman, and W. Carleson for red fox samples.

## Data Availability

We have deposited the primary data underlying these analyses as follows:

- Y chromosome variable call format (.vcf) file: Dryad
- Sample information, primers, and flanking sequence uploaded as online Supplementary Material

