## Supplementary Information 1 for "Sequencing red fox Y chromosome fragments to develop phylogenetically informative SNP markers and glimpse male-specific trans-Pacific phylogeography"

**Supplementary Information 1. Fasta sequences (and Genbank Nos or references) used to synthesize 2x overlapping 80 bp RNA baits for the Canine Y chromosome.**

**>Fragment 03 (**Natanaelsson et al. 2006; Ding *et al.* 2012)

TTGTCTAAAGTTAAAGTCACAGTACATTAACAAACAAATGATCCTCAGTCACAAAATTATAAATGCAGTTTGGGGTGAGAATAAGCAGGAGGGGAGTTAATCCAAACTACAGCAATTAAGTCTTCAAACCAACTTGTGAATAAGATTGTTGATTGTTTCTTTCTTCTTGTGACTTTGAAGCTCCCTTGAATTTTCAGGAGAATCACCAGCTGGTAAAATGTACTGCAAGGTCTTGAAGCTATAAAATACAGTCTGGCTAAGTTACATGTGGTTTCCATATTAACTTAAGGGTGGCTTTGTCGCACAGCAGATTCAGTTGCCCCACCAGTCAACCCCCTGGGAGTTGTAATTTCCTCCATATCCATCACTACTGTAGAAGCCTCCATAACCACCTATCAAAAAATGTTTGAAAACAGAAATTAGTTTAGCCATTTGCCTCTTCTTTCTCTGATAAAAACCTAATTAAAAAAATTCATAGTACTGGATTTTGTATTCTTTGGAAATAAAATTTCTCCGATGAATTATGGTTTAAGAAAAAAGTAAAAAGCACTAAAATACAAAAGGTATTACCTCCACCAAATCCTCTGCTGCTGCTGTGACCACCTCCACCACTACGGCTGTTGCTTGCACGACTACTGCTAAAGCTAGAACTGCTTGAGCCACTACTTTGTCGA

**>Fragment 11 (**Natanaelsson et al. 2006; Ding et al. 2012)

TTATGTTACTTATTGATAGTAATAACTGGCATTTTAACAGCCCTGTTCAATATGATTTTAGATGATGAGCTTTTATCACTAAAATCAAATAATAGAGGATACTTGCTTAGGAAATTGAAAGTCCAGACTAACCCTTGAAATAGTATGACGCTGAGATAAACTGATTAAAACCCTTGGGTGTTTTAATATTGTATAACCTCATCAAAATATGTTATGATAAGATCCACAAGATTCTTCTAGGGTAGGTGTTATACTTCAGATAAATATATATCTTTTCAGTATTTGAAAACAGGTGTTAATAGTTATTATTAGTTGTTCAGCTGGAGCTTTACAAATCATTGTGTTTTCTGTAGTAAGCAATATCCTTATTGTCATCTGGCATAATAATTCTCATGAATATACTGTTCATACTTACTTTCCTTAATATTGAAGGCTTAAAACTGACCACAATTACTCATTAGTTTCCAGCTCTTAAAAATAACTAATTATTGGGACGCCTGTATGACTCCATCAGTTAAGCATCTGACTATCTCAACTCAGGGCATAATCTCAGGGGTCTGAGTTCAAGCCGCATTTGGGGGGGCATGGAGCCATTTTACAAAAATAATGGTAATCTGAATATGTATTATGACAGTGATTTTGTTTTTCCTTTTATTATTAGCTTTCATTAAATATTAAGCTAATTAAGCTATGATATATGCTCTGAAGGTTCAAAAATGAATAAAACATAGTCACCACCTTTATGGAGCTAGTTTGTGAGGGAAGAAAAGTATTTAAATACAAAACACTGTGAATACTATCATAGAGAAGTAAAAAGAATCATTACTTTTACCTGGTTTTCTCCTGCTATATTGGTTAATTAAAAATAGGCTAACCTTTATTAAACTGAATGAAACTTAAGTGGTTGTAATGAGTATTGAATTCCCATGGATCAATCTTTAAACATATAATATGGCTCTTGTTTTTGTGGTATAAGTCAACCTGGTGTCTCAGACAATTAAGAAGTATTTTTAAGTACTTCCATTTGCAAAAATCACTGTCCATAATTTCATGTAAAGTAGGTAGAATTCGTAGTTATTCTAAAGTACAATAAACAACAAATCTCCATTCAGTGCTTTTTCTCTTATATG

**>Fragment 12 (**Natanaelsson et al. 2006; Ding et al. 2012)

GTCGACTTTGAGAGAAGCTCTAATAACAGCAGGAAAGGAGGTCATATGGCATGGGCGGACAAATGATGAGCCAGCACATTATTGTAGCATTTGTGAGGTAACTGTTCTCGCCAACAGTTTTTTGTTGTGGGATCTCTAGTTTGGCGATGCCTTCTAAATTACAAATTTACCAGTACTTTTATATAATGCATTTTTGATTGGTTATTACAGCAATTAGTTTTAATTTCCATTTGATGGTGTAATATTTAATTTAATTCAGAAGTATTGTCTACAGAAGCATTCATGTAGTACCTTCTTACTAGATGATAATTATCTCAGATACTGAACTAGAAAAATTTAAAATAAGTATAGAGATACCAGATAACTGGATCTGTTTCAGTATAAATTCAGGCATATCATTCATAATATCATTCATTAATACAGTTGTTGGTTTTCTCCTCTCGCCAAGGCTTGCAAGAAGTACAGAAATAAAGAGGTAGTATCTCCTTTGGAGGAGCTTCCTTCCACTAGGAGAAGACAGAATGTAATGGATTCTGTCCATGTAATGGTTATGGATGCACATGAACTATGACATTTGAAGCAATGAGGAAAATAATCTAATACTGGCTTAGAGTTTATAGGCTGTTTCTAGAGTTGAGTCATAAAGTCTGTAAGAGGAGATGGTTTGTAGCTTGCAATGATCTTGAATTAAAGTTATGGAATTTCATCTTAACTCTGAGGTCGCTGAAGCCATAAA

**>Fragment 16 (**Natanaelsson et al. 2006; Ding et al. 2012)

TGACCTCTCCGATGACAAAAGTAAGTCCTCCGGTTATGTATCCATGAAGAGCGTTTAGAATCTCAAGATTGGAAATTGTTTTGTACTTTTGCAACCCTACTCATTCTTTTTGTTGTTTATGTTGTCCTAAAGTCTTAGTTATATATATAACATTCTGATACAGCATTCTGAAATGATTTTTGCTAGTTTTTGCAATTAGATTTTAGTTACTGAGGAATCTCAAAAGGTTAAATTTACAAAATAGAGAAGTTACTTTATTTCTAAAATTCTGTTTTTAGGATTCAATTTTCAGTGATCTTAATTTATTTTCAAACTTTTTTCTCTGGAAGAGCTAACACTCTACTTCTAACCTTAGAAGAGTCTATTAGAGATTTAAATGTATTTATTCGAACATAGTATTTTCAAAGACCTAGTAAGTTTTCTTCAGTATTTGCAACAAGGTGACACTTTTGATGTATTACTTTATTTCAGGTGGAAATTGCAGCTACATGAGCTTACTAAACTTCCTGCCTTTGTGCGTGTGGTATCAGCAGGAAACCTTCTAAGCCACCTTGGTCATACTGTGCTGGGCATGAACACAGTTCAGCTATACATGAAAGTTCCAGGGAGTCGAACCCCAGGTTATTTTTGTGAACTAACATATCTTGTATAACTGAAAGAAGAGTTAATCAAATATAAAACACACATACAACCCGGTTTCCCATTTGTAATTAAAAAAGCCAAAGGACTTCTTAATGCTGTCTCATATGTA

**>Fragment 20 (**Natanaelsson et al. 2006; Ding et al. 2012)

TTGTTTAGGTTATACTCGTGATTCGTAAAATACTCTCTTGGAAATACTGTAATACAAATTTTGCTTTGTATTTTAGGTTGGCTTTTTTTTTTTTTAAGAATCATTTTGAAGACTAACAAAATTTTTATTACTTATTAATAGTTGCTCTCACACAGAAAATGTTGAGTCTTGACTCTGATAGCTCTAAAATTACAACTTCACTACTACAATATCATACATCAGGTCAGAGTGATAAGCTTCTGTTTCTCACAGAAATTCATGATGCTTATTTTTATTAAATCTTCAAGAGTGGTACATAATCAAGTATCTGCCAGTTGACTAATACAGTGTATAACCGAGGTATAACTTTGCTAGTTTGTCAGTATTCATTCTACTCTCCTTTTACACATTTTCTCTTCCATTAGTTATATTTCATCCTAGTAGTATTTAATGTTTTTCAGCTTCTTTAAAAATATTCTCTGTAGTTTGTCCTACAAAGACTATGTTTAAAGATTAATCTGTCAGTCACTTAACATTCAGCCTTGATTCTTCAGTATCACTTAAATAATAGTAACATCTGTTGACCTGTTAAGAACCAAGAAAACTGCCTAAAATTATTTACCTTTTTATTTAATTGACTATTGGTATTTGTTTCAACATCTTTATTGAGTGGCTGTTCTCAAAAGTCTTAATTGTCTAGAATCATTTTTATTTTCTTTAGATATATTGATTTTACTCAGTGCAGTGAAACACAATTTAATTAACTGTTTACCAGTTTTCTTGCTTGACCAGAGACAGATTTTGATTTGTCTCATTTTTTCCTCTCCTTCTTTCCCTCCCTTCCTTCCTTTTCTTCCTTTCTTTTCTTCTATTCCTTCCCTTCCCTTTTCCTTTCTCTTTCCCTTTGCCCGACATACTGCATTTACATTTCTAATTTTTCTGTCCATTGTTTTACGATAAAATAGATGTATTTTATGAGCACTTTGCTAATTTCAGTGTACAATAATTGCTTAGCACAGCAAGAACATCCTTGTATATTAAGTACAATATTTCTGTGAAATTGAGAAAATAATTTACACTACTACTTAAAGTTCATTTTTCTTGCTTTCACACAATGGATATCCACTGGTCTTTTTTTAAATGAAATGTTGAAGTATGGTGATATGTAGATCAGTTGAAATGCTGTTGACTTATAACTGCCTTATTACAATAGTCACCAACCAGCAGTGACAACTTATGATATTTTATTAGATTTTTCTTATCTAAGAATCAGCTTAATTTTTATTTTGTTGTAAATGAAAACATCTATCTACAAAACATAAGTCAATGGCATATTTTGTTCTATGAAAGCATTACTTAAAAAAAAAAGAAAGCATTACTTAGATGAATAGATGTTTGGCCTATACGCCATAGGTTGCCATCTTCTGATTTAAAATCTTGAAAATCTGGGCAGCCCCGGTGGCTTAGCAATTTGGCACCACCTTCAGCCTGGAGCGTGATCCTGGAGACCGGTGATCGAGTCCCACCTCAGGCTCCCTGCATGGAGCCTACTTCTCCCTCTGCCTGTGTCTCTGCCTCTCTCTCCTGTGTCTCTCATGAATAAATAAATAAAATCTAAAAAAAAGTAAAAAAGTAAAATCTTGAAAATCTAAAAATTTTTAACCAGCACACATCATTGTCTCTCTTGCATTGACCCAGATTATGTAATGGGTTATAATGCTAGTATTTAGCAGTCTGTGAGTTCCAGCCAACTGTCTATTAAGGCAGATAATAGCAGCACACAGCCACTCCCTGTCTATATGTGATGTCTGTAGCCTGGGTATCACCCACTTTTATTTTTATTTCTTGGATCAGCTATAGTTTGACAAGCCAGAGTTTCTTCACATTTTCACATTTTC

**>Fragment 21 (**Natanaelsson et al. 2006; Ding et al. 2012)

**T**TATGCTTTTTGTCATCCTGCAATGTTGCCAGAAAAATATACAGAGAGATTGGGGGCAAGGGAGGGGATAGAATGCAGGAGAGGGAGGGAAAGAAAAGAAAAAAGCAAAAGTTATAAGCATGGTACAAACCTTATTTCACCCAAACACTACCAATATAAAACTAGAGAAAACTTCCCATAAAAAAAGGAAAAAAAAGCCAACTAAGATAGATAGACCATGCATTTTTTAAATAAAGTTTACAAACAATGGTTGATTTAAAGATATTAAGTATAGTTATGAAATAAACTATTAAAAGTTTTAAATATTAGAAGCAATATTCGGGATGCCTGGGTGGCTCAGCGGTTGAGCGCCTCCCTTCGGCTGAGGGCATGATCCTAGAGTCCCAAGACTGAGTCCCACATTGGGCTCCCTGCAGGGAGCCTGCTTCTCCCTCTGCCTATGTCTCTGCCTCTCTGTGCCTCTCACGAATAAATAAATGAAATCTTAAAAAAAAAATAGAAGTAATATTCTGTTCTCTGCAATATAAAATTTACCGAATATCTGAGAGGGTAACTTCAGGTGAATTAGTATAACCTAATATTACTCCAAAACTCAAACTTTTCATATATCACTGTTTTCTAACTATCTTTGACCTAAGTAATTACAAAGTAATTACAATACCAGTAAGGTAATCTGTCTTATCTGTCTTATTTTAAAGCAGTTAAACTCTCAAAATAGTTCATACTCAGAAGAGTAAGTAAGGTGCTGTATGTTTAACAGTTACAATATACCTCCTTATAACTTTAAGAATTAGTTCTTGGGGCACCTGGGTAGCTCACTCAGTTAAGTATCTGACTCTTGGTTTCAGCTCAGGTCCTGATCTCAGAGTGCTAAGATGGATCTCCAATCAGGCCCCATCCCTAAGGGCGGTGTCTGCTCCAAATAATTTCTCCCTCTGCCCCTCCCCCTCCTGTGCATGCATGTATCCCATATTCTCTAAAGTATTCTTATTCACTCTGAAGTAAATAAAATCTTAAAAAAAAGAAGAGAAAGGAAAAATATTCTTAAACATAATCGAGATTTCCCATCAAGTGTGTCCTTCTCCCTTGCTTAAGCTACCAGTTTGTTGATAAGAAACATTCAGAAAAGAGATTATTACATTATTTTCCTGACAAATTATCAATGTTTAGAAATCTGACCTCTTTTATCATGAGAAAAAATTAGCCGTGAGACAACAAAAGGGATCATTATCTTGCCAGTAAGAAAACTCTTTAAGTATTATTTAAGCTACCATAAAACACTTTCTGTGAAATCAAATATCCTGAGTGAAGACCCTCCGAAATTGATATAAAATGTATACTGATTTTAAAATAACCATACATGTGAAGTATTAGAATTTTAGCAAATTTTTTATATTTCATAAACATATCATTTGGCAGTATCATTATAAACTGAAAATAGCAGGAGACTGACTTGCTACTGATCATGGCAATTTTAGACTCACTATTAGAAATATATAAGCTTGAATTATTAATAAGGTATAAGGATGCTTAATTTTAGAATTCTTCCTATTTCTAATTATAATTAAGTCATTTTTTTTCCTGTAAAGCACACCTGAAAAAGAATGTGCATTCACACTTATCAATTTTTAAAATTAAAAAAGAATTATTTACTCCCTATCCTTTGAATTCGTACCTGTTGTTATTGAATATACTGGTTTCGTGGCCTATTTCACTTAGTGTTATTCTACATAACATAAAATGAAATATTTCAAATAATAGCCATTTAAAATGAAATTTCTAGATTTTGGAAATTTAGCATACTTTGTCCTTTTTAATAAATGTCTTCTCACAATTATTTCAAAAGAAAATCTGTTCAGAAACCTACTTTTCACAAAAGTCATTCAGAACACCCCAG

**>Fragment 24 (**Natanaelsson et al. 2006; Ding et al. 2012)

ATTCTTTAAAAAAGGATAAAATGTAAATTATTTAAAATACCTAAATGGGGAGTGCCTAGGTGGCTTAGTTGGTTAAGCATCTATCTCTTGAGTTCACCTCAGGGTGACGAGTTCAAGTCCTGCTTAATTTATTTAAATAAAAGTCAATAAAATTTTAAATTTTATTTAAGTCAATTTTTATTTAAATCAGATATTTAAGGTGTTTCATGCTGTTTTTCACATTTAATTTGATAATTGCCTCTTTTGAAATTTTCATGAATTAAGGAATGTTTGTTCTGAGTATTGTAAATTTATAAAGGAAAATGCGGGTTATCAACATTTAGGTTTTTTGTTTTATATTTTGGTTCTTGTCCTTAAGGAAATAAAAATACAAATGATGTATGAATAGAAACCTGTGTGCTTAAACCTGAATCTTATGGCATTGAACAAATTTGGTAATGATCTCCATTGAGTTCATTAATATGCAGTTGTTAGCATTTATAAGAGGAAAGTTCATTTAACAGCAAATTATAATATAGATTATAATTACTATGATTTTTGTGTATGTAAGCACATACGCACACATGTATAGTAGCTTTTTAGGTTTCTTTCTCTATACATCATAACATAAATATAGTTAACTTGAATATTAATGGAATACTTTATATATATGTATACTTTTACTAATCTCTCTCTACCTATACTCAAAATACATACAATTTTTTCCTTCCTTTCAGGTGTTACTCAAGTATTGGGAAAGTTCAGGATGCCTTTATATCTTATAGGCAATCTATTGATAAATCAGAAGCTAGTGCAGATACATGGTGTTCAATAGGGTAAGCTTTCTTTATATAAAATTAATAGTATTTTCATAGATAACAAGCTCAACAAGCATATGTAATATACTGTTTATTAGTATTATACTCAGTTTGTTTCAGACAAATTTTTTTTCTTTTCAGTGTGCTGTATCAGCAGCAAAATCAACCTATGGATGCTTTACAGGCCTATATATGTGCTGTACAATTGGATCATGGGCATGCTGCAGCCTGGATGGACCTAGGCACTCTCTATGAATCCTGCAACCAACCTCAGGATGCCATTAAATGCTACTTAAATGCAACTAGAAGCAAAAGTTGTAGTAATACCTCTACACTTGCAGGAAGAATTAAATATTTACAGGTAAAAATTTGAAATAGCTTATTCTCAAGAACTTTTCCATATGATAACTGTCATGAGGAGGGTGGCTGGAAATTGTGTCTTTTTTGTTTTCATGTCAGTAACTTTTTTCCATAGTTTTTAATATTTTGTGATATTTTACTTTTCATTTAAAGTAAAAGACGTAGTATTTAGTGTTCTTTTTTTGCACATTTACTTTGTTGAGATACATTAATTTGCATGATTTTTCTGTGTGCTATCTACATTTTTAAGTTACAGCATTTTAAAGGAAACATTGTTTTCCACTCTTGACTGAAATTTCATAAGATCTTTAAGCTTTCTGAAAAACAGAAATTACCTTCGTTGTGACTCACATGAGTATATTCTAGTTGCCTTCTTTGTATTCAGGATATATCAAAACCCCCGTATTTAGACTCTATTTTCTTTGAAAAATAAATACCTGTAACATTGAGGAAGATTTAGTGGACTTGCTCTTACTCAAGTAGGCCTGTGTTAAATTTGCTTTTTACATAATTTTTCCTAGGCTCAGTTGTGTAACCTTCCACAAGGTAGTCTACAGAATAAAACTAAATTACTTCCTAGTATTGAGGAGGCGTGGAGCCTACCAATACCCGCAGAGCTTACCTCCAGGCAGGGTGCCATGAACACAGCACAGCAGG

**>Fragment 27 (**Natanaelsson et al. 2006; Ding et al. 2012)

CTGAAACTGTTATTTTAGATAAATGAATATATCATAGTTTGTTTTTTTAAACATAATTTCTATTACTTATAAAAATCAATGTATTATTTCAGTTTACTAATTCTGAACAGTAGTCAACCATTCCCATTTTGTGACAGACTTTCCCATACTAACCTGCATGCCTTCAGAACTTCCGCAGAACTTGAGCATATGGACACAGACATTGATGGTATAATATGAGTACCAGGTTTTTGGCTAATTATAGCCAGCTCTGCTTTTATAGAGCTCTTTGAGTTCTCTAGACTTTCTCCATTAACCGTATATAACCTACTGTGAGGACTGTTAAGGCTGGGAACACTGTGTGTGGCTGTTTGTTCAGTGGATTTTGGAGAAGGTGTTGCTGTAGAAATGACTGAAGAGGGTGAAGAGGCAACAGAATTATTAGTCTTTGTACAAATAGTTGGGTGAAAATGATTGACTTTGTCACAGGCTTCCCTACCCCCAATGTCATTGGCTTTTCCAATCAACAAGGCAGAGATCTGAGGATTGTCTGAAATCTGTACATTTGGTGAAGTGGAATCTCCATGTTTAGGACTGCAAACAGGTTCTGCCACCAACTGCTGAACATGATTAGAAAGTCCTTTAACACCAGCACAGCCATTAGGTGTATCTTCAACATGCCTGCTTGTTTCAGGCACCAAGGATCGATTTCCTGAAAGTTTGCTCTCTTTGGTAAAGGTAATGCCTTGTTGTCCACCTGAGGTAGGAATATGAGAGGAGAGGTGATTGAGAGCAGCTCCCTGTGTTACTGAATGGCTAGGCAGAGTTCGAAGTCCAGCCTGTTTCTGAATAGCAGTGCTAGCTGCCTGGAGATACTGGGCAGGCCCAGTGGAAAAGGAAGGATGTTCATCATTAGGTCCTGCCAAATGTGAACTCTGACCTTTGTGAAGCCCCTTAAAAAAAAAAGAAAGAAAGAAAGAAAGAAAGAAAGAAAAAGAAAGAAATTACCAAGTTTTGTTTTAGTGTATTATTATACAAACTCCGAATACATATACAGAAGGACACGGAGTATTAAAAATGTTTAATGATCTATGAAAAAGCAATTAAATGTGAAACATATTTTAAACCAAAACCAGAACTTATTTTCAAAAGTTCCCATAATCCAATATTAACATGGTTAGTATATTCAGAAAAATTATGCTGAGCTGTGCTACACAATTTGATTCTACATATTAGTATGTCAAAATATTTAATATAAGCCTTTAAGAATTTTTAATGTTCAATTCTGATACATACATTTTCACCTTCTGAAAAATATATTTATAAATTTTGATATGGGGGAAGATACATACATAGTCATTACCAAGTGAAACCATGAAACAACCCAAATATAAATAGCACAAGTCACTTATTTAAAAACTATTTATTAATTTAATAACTTAAATGGATTTCCATATTCTCACTCCAACCCCTCATAACTGACAGACATTTTAAGTGCAGTTCCCCACTAATTACTCAAGAATTGGCAGTTTTACCAAAATTCACATGGATTTGAAGAATATATTAAGGTTAGAGCATTTTTATGGGATAGTTTATTGAAATCATTAAGTTTTGAACCAAAAGGAAGTTATTTCATGGAATTAAAATATGCAATGAATTAAACTAATTTTACCTGAAATTGTCCTTGGCATGGTAAACTGTTCTAAAAAGATTATCATTTTATTTTATGAAAGTAATTAAGCAACATTTATTAAGTAATATGGAATAAGGAAACAATAATACGTTGCAAGCTAACATAACACTAGAATGTGAAAACTATTAACTTCCCTGATGTTAGAAATATAATTAGACATTAATACATCTTTAATATAAAACTCTATATGGAATTAAACCTCAAAACTGTAGAGTCTATCTTAGAATTGCTTCTTTCTGAATTGGTAACTAATTTAGCAACCAAGCAAAAGCGTTTGCTGTCATAGAAATAGAAATAAGTTATATATACA

**>Fragment 28 (**Natanaelsson et al. 2006; Ding et al. 2012)

GAACAGAAGTGTTGAAGAATTGGGAGTGATCACTCAGTCAAGTTCATTTATTTCCTCATTCTATACACAGGGGATATGAGTCAAGAGAAGTAAAGTGGTTTCCTTAAGGGTACTGAGTGTCACAGAAAGGGCTGGAACTCAAGTTTATTTGGTGCTGCCTGTATTAGAAGCCAGCCTGCCTTAAAGGAATCTACATAAATTGTTTTGGGGGAAAAGTTCTTGGTGATTGGACTTAGATATTAGTATTCGGGAACATTGACATTATGTTTTGACTTTTAAAATGGTATGTGATAATGCACTTTTCAGAAAATTTTTCTTATTTTCCTAGCATTTAGGCAAGTGTGTCTGATAATCTGAAATTTGGTTTTGTTAAAATTTTTTTCTGTTGCTATTCACATCAACTCTGTGTCCTTGTTTCCAGTTATTACTGACTCTTTTTTATATTCCTTCCAACGTGGGTTTTGTGTGCTCTCTAAAATTGCTTTCTCTGTTTTCTTTTACTGGTGTTTCTTCTATCTCTTAACCCACAAATTAGTCATTCTTTAGATTTGTTTCTCCCTCACCTCTTATATCCATCTTGTGTTCTTAAGCCTCTCTTTCATATGGATAAATTGCAAATTAAGATTTCTAACTCTTAAGGTTCTGAACTTGTTTCATAATTCAGTTCTGTTCACACCAAAACTCTAAATCAGCATCTTTTAAAAGCTCGTATATCGTGTTCTTTCTCAAAATAAATACTTTATTCTTTCTCACAGTAAGTTACCTTTTTATATGTTTTACAATTAAATCTCCTTTAGTATCTAGTGAAGTCATTGAGTCTTTATCAATCATTTATTATTATTCTTGTATATAATTTTCATTATTTCCACTATAATTGTGGCTCAAGAGTTTGTATCAGCCTTGTGATTTTCTGTCTCAAATTTATTTTCATTGAGTCTTTAATTTTTCACTTTACACATTGTTAGAAGATTGTTTCTGTACATTGGACCCTTGCCTTTTTTGAAGGATAGATTCAAGTTGATCACAGATTTAAGTTTTCCATTTGTAGAGTAAGTTTTTAGATGATATGCTTTTTGTCCTCTTTCCTACTATAATCAGTGGTTCCCAACCATTTGGGTTACCTGAATTAGCAAAATTTGAATAAAAAATAGTAGTGGTCACCTTAAAGTGGCTATGGAATCATGTGGTTCCTGAGATCGAGCCCCATATCAGGCTCCCTGCCTAGCGGGAACCTGCTTCTCTCCCTCTCTCTTCCTTCTGCTTGTGTTCCCTCTCATGAGCTCTCTCTCTCTCTGAAATAAGTAAATAAATAAAATGTTTAAAAAACAATTTTGCTATTTTGTATCCTCCAATTAGAACTGTATATACAAAGAATTCCATTGAAATACATATTGAAATAATTCCTGTCTTAAGAAAATAAATTTAAAACCCAGACTACTTGTTGGCTGTGATTTGGGTAGTAAAGTTTTCCTTTCTTTAAATCATTGTAAGTTTTGTGTCTTTGTTGGTTAGATGGTTTTTTGGTAAAGTAGCCATATACTGTCCTTTACTTTTAACATTTCTTAATATAACTGCTTTTTACAAAGTAGGATTAGGGATGTCTGAGTGACTCAGTGGTTGAGTGTCTGTCTGCCTTTGGCTCAGGGTGTGATCCTGGGGTTCTGGGATCAAGTTTCTGCATCAGGCTCCCTGTAGGGAGGAGACTGAGAGGCCTATGTCTCTGCCTCTGTGTCTATCATGAATAAACAAAATTCTTTAAAAAAGGATTAAAATGTAAATTATTTAAAATACCTAAATGGGGAGTGCCTAGGGGCTTAGTGGTTAAGCATCTATCTCTTGAGTTCACCTCAGGGTGACGAGTTCAAGT

**>Fragment 29 (**Natanaelsson et al. 2006; Ding et al. 2012)

ATAGATTCCCCCCTCCTACATATCTGAATTTAGCTTTTTAAAAAATAGCTTTACTAAGGTATGATTCACATGTCATACTTTATAATTTCTTCTATAGTCAAAGTGTAACCAAAAAAAGAAAACACAGTGTAGTTTTATTTATTTTTTATTTTTTTCACAATGTAGTTTTAAAATGTACAGTTCATTGATTTTTAGTATATTCCCAGATATGTGCTACTATCACCACAGTCAATTTTCCAGCCATTTTAGCACTTCAAAAAGAAATCTAATATCAGGTTTTTAATGTAGTGGTTTTATTCTGTTTATACTAAACCATAAAATCCAATCCTACATTAAATAATAAACTATCTTTTTCCTTTCTTGCATTTACTTATAAATGTCCTTATTAGACACTACATAAGGGATGCCTGGGTGGCACAGTGGTTAGGCTCAGGGTGCTTTTGGCTCAGGGTGTGATCTTGGGATCATCAGGCTTCCTGTGAGGATTTTGCTTCTCCCTNNNNNNNNNNNNNNNNNNNNNNNNNNNNNNNNNNNNNNNNNATAAATAAATAAATAAATACTTTTTAAAAAGACCCTAAATAAAACCTGGATAAAGTAAATGCTTGAGTAAATATATGAAATATACTGTAATATTTTTATTATATAACATTTGAAATTTGAACACTAATATGTTGAATTTTAAAAAGATATTCCTAATTTGAGTTTGAAATAAAATAGAAAATAAATGTCTGTTAAGTACTTTTGAACTACCTTGACTATTTTATTTTAGTATAAGTTATGGATATCCAGGTGGAAAATAATCATGCTTAAAGATTGTAAATCTATTCATGTAACTAGTAGTTAAATATTCTGTATTAGCTGTATCTTAACCTTGCTTAGACAGGAGTCACTTAGATTATTTTATTAGGATTATGAGTCAATCATTAAATAGTGTCACTGACAAAGTTATATGATACAGAGCACATTTTAAAATTTTTATGTGATTTGTACTTGCAGCCACCTGGTGAGTAATGGAAATAATATTTAATGTAAACTTATGAAAATTAAATTGGTAT

**>Fragment 30 (**Natanaelsson et al. 2006; Ding et al. 2012)

CTTGAAAGAAGTGCCAGAAGAGAGAAAAAGGTAAAACTCCAATTCAGAAACACATACACTTACACAGTAGATGTATACAGTATCTAGTCTTCCTTCCTTCTACCCCTCGCCAATGTTATTTCACCTCTGAAATAATACCTGAAACTCTGCAAATTTTTATTTGCATATGTAAAAATGATTATAAACTCAGCAGTGCTGCCAGGTAGGAAAATAAAATGTTAACAAGAGGTAGCATTCATAAAACACGGAATTAAAGAAGTTACACAAGGTTAGGCCACTTAAATCTAGTGACACTGAAGTCCAGTGCCAAGAGCCTTTATCATCCATCTCTTTTCACAATGAGGGAAAAACTGGGCTCAGAAAGGCAAATTTTACCTATGTTCTCCTGCAATCACAGGTACAGAATTACTTGCCCAGATAGTGGAATCGTAAAAACTGACACTGCTATTAATATTATCTCACATAATGGATAACCAAATAATCAATGAAATGGTCATTCCATGTCATGTTTTCTCCATTACCCTGAAATTTTAGTCCTTTCCATTTTCAGAAAGCTAAGGAACAAAAGAATAAAGGACAATGTAAAGGTATAAGGACTTACAGGTAAAAAAGTCAGTAAGAATCTAATTTTTAAAAAAAGATTTTATTTATTTATTCATGAGAGACANNNNNNNNNNGGCAGAGACATAGGCAGAGGGAGAAGGAGGTTCCGTGCAGGGAGCCCGACATGGAACTCGATCATGGGTCTCCAGGATCACACACCCTGAGCCGAAAGCGGCACTAAACCGCTGAGCCACCCGGGCCACCCCAAGAATCTAATTTTAAAAGCAACAAACATAAATAGAAAATAAAGATACTTAGGAAAAGCATCTTAAGGAAAAAATAACGTTCAAGACTTAACAAACTATGCCTTACCTCTCTCTAGATCTTAATTAATCTGAAGAGAACTTTTCAATATACAGATGGCTTAAAAGTATCAGCAGATAATTTTTGCTTGATTTTGGGAATCC

**>Fragment 31 (**Natanaelsson et al. 2006; Ding et al. 2012)

AGCAGGACCTATATGAAATATGCTCTGGAAAAAAAATAACCCACTAAGAATATAGTAGGTTTAAGAGAATGAGATGTATTTAAGTATGTATACATGAGTGGTATACACATTCCAGTCAGAGTGTCAGCAACTAAAAGCATAATTAAGATAGAGGTTTGAAACTCAAAAGATCTGAAAGAAGCTGGAATAAAAGACAGTGCTGACTCCCAATGCCTATTTTGTCAAATGGATTAACTATTAATGGGATTCTTATCTAGTCCACTTCTTATATCTTTAAACTGTAGACACATTTATGATTTTTTTAAGTAGTTCATAAGATTCTTAAATATATATTTGTCATATCTCCACATAAAGATAATACTATATAATATTTCATTTGGAGGGACCACAGTTGCTTCCAAATTATGGCAGTTATGAATGAGGCTATGTAATTCCTTTGGCCTTTCATAAATGCCCTTCTTTATCAGTTTGAGATAATTTCCCTGTATTCCTAATTCTTTTATCGTTTTTATCATGAAACGATACTGAATTCTATGAAATGGTTTTGAATTTTGTCAGAAGATGTTGAATTCTTACATTTTTTTTTTTCCTGCATCAATAGAACTGACCATATATTCCGTTTTAGTTCATTCCTTTACCCTAGTCTCTTGGAACCTTCATTTTACCCTATATATCTTTGAACTTTTGTATTCTAAATACTTCCTACAAACAGAATCATGCAGTATTTGTCCTTTTGTGCCTTACTTGTTTCATTTACATTACCATCTGTGTCATCTCTTGTCTAGCAGGTATCCAAAA

**>Fragment B (**Natanaelsson et al. 2006; Ding et al. 2012)

CTGATAGGAAGCATGTTGTCTTGCACTTGATGCACTGACGCTCATCATCTGGAAGCAGCTCAAAAGCCTCTCGTTCAGCCTCCGTGATGCCCTGGGGGTGTTCAACCCACATATACTGTAAAGATTAGGAGCAGGGTTGTGAACACCCCACCTCAGCCAACACTGAGCCCTGCTGTTGGCTTAACCTTGTAAGGGTAGCTGTAGAGAAATCGGAAGAAAAAAATATAAACACATACAGTTTCTGTCTTTCCACTTCAAAATGTATGAGTCAGAGATGAGGCACAGAATGACTACTCAATTTTTATCTGTCTCCCTTCCACACATCTGTTATTTTTTTTTTTTTCACACATCTGTTTTGAATCCCTAATTTGTACTGCCCACCACCCACCTTCTCCAAAAGAGCTTTTCGTAAACGTCGCTCCTCTTGCACCATAATGAACATCTCCTTGTGCACAGCCACAGCCAAATTTAGGTCCAACTTCTCTGGAAATGCAGCCATCTTGCAGATGAGCTCTTCATGGGAAAAAACACAGTAGCGCCGGAGCCGGCGGTAGTGTTCAATGCACTGGCGTCCAGCAGGTAGCTGTAAGTAGTTCAAGACTGTGTATGGCCATCTGCAGCCCACAGTTTTTAGCTTCTCCAAAATGACCATATTCTGTCCACATTCTTTATCCCTTTCTATCTATAAGCCTACCACACTCCAGACTCACCCAGTCAGCAGTGCAAAAGTTGACAGCCTCAGCAAAGTTGTAGCCTTGATTAAAGCCACTGTGGTAAGCACGAGGAAAT

**>Fragment G (**Natanaelsson et al. 2006; Ding et al. 2012)

CCTGAGCCAAAGTCCCACGCTCTACCCCTGAGGCCCCCGGGTGGACCTTCACTCTNNNNNNNNNNNNNNNNNNNNNNNNNNNNNNNNNNNNNNNNNNNNNNNNNNNNNNNNNNNNNNNNNNNNNNNNNNTCACAAAAATAAAGTAGTGGGAAATACGGATAAAACTGTGCTATAGATAAAAATGTACTTCCATGGAAAGCATTCTTTGAAGCTTTCGTTTGAAGGAATCCCTTTGCAAAGGACAGGTTCATTGGTCTCTCAGCTACTTTGCCTGTCTTTGTAATCCCCAGATAATTAAGGACTACATATGTAAATTATATCTTCTTTGGTACTGTATATTATCTCTCCTTCCTGCTGTGGAAATCCGGTTTTGTCCATATCTGCAGTTTAATTCCTCTCCTCTACAGGTATTTGCGCACTTTCCCTTCCACTTTAAACCCATCCAGTAACATCCATCCAGTGCAAGAAAATCTTTCTTTTTTTTTTTCTTTTAAGATTGTATTTATTTATCCATNNNNNNNNNNNNNNNNNNNNNNNNNNNNNNNNNNNNNNNNNCATCCGCACAGGGAGAAGCAGGCTCCATGCAGGAAGCCACCCGGGAGACTCTGGGATCACAGCCCTGTGCCAAAGGCAGACACTCAACCGCTGAGCCACCCAGGAGTCCCGAGAAAATCTTTATGTAACTCGTGTTCATTTTTATATCTGTAGGACACCCGGACTTTTGCCTCTTTTACTGAAAATTATCTTCTAACAATGTAATTGCTTCCATTGCTGACTGACTTACACACGCACAAAGGTAGAGGAAAAAAATATTGAGAATATGAACTATTCAGATTATGGGGCAGTGGGATAGGAGGTGGGAGAGCCAAGCCTGGCTAATCTAATGTAGATGTCAATGAAAACTTAAATCAAAGGCATTACCAAGTTTATCATACAGGCTAAGGTAGAAAACAGAGTTTGCAAAGAAAATCATCACTTTTTAAAATAAAGATAAATTGATGGTCAAAGTACATACCGTCTATCAAAACCAACTTTCCAATACACATATTCCAAAGTTTTTCAAATACTAAAGCTCAAACACTAAGATAGTACATTTTATTTTTTATTTTATTTTATTTTTTTAAATAGTACATTTTAGAGCTCAATGCCATGTTTT

**>Fragment K (**Natanaelsson et al. 2006; Ding et al. 2012)

TATATATCCCAAAGTATATCCCCCCATGTGTACAAAGGCATAGATGTGCTATGTGCCAAAATATTTGTTTTTGATATTCACTTATTAATTTGAATACTCTCTCATACTAACAAGATTTTCCCAAGGAGACACAATATTCCAGTATTTAGTGAATTATTTTCTAATAGAATAATATCAATTAAAAATAATCCTTTAAAATGTAAAATCTGTTAATGGGTTATTAAAATACCTGTAAGTGGACCAACATTCCAAGCAATATTGTTGCACCAGCCAACAGCCTGAACCCAATGCACAGTGCCTGCATTTATCCAGACCAAATCTCCAGGTCTTTGAATAAATCTATATACGGGGACATTTGCTTCATAAAGATCTTCAAGGTTGGGCCACCAAGAACTCATCAGGAAATTCAAATTATTTCTGGGGGGAAGGGGGGTAAAGGGGGGGAAGGAGTAAATAACATAATGAATTAAGTTTTTTAAAATGTCCCATAAAAGATGAGACAATTTCCAAATGTTAAAAGATACTTTCATTATGCATTTCATTAAAACGTAATAGTACTGTAATACTGTACTATAACAAGAACTGCATCTTTCAAAAGTGTAAATATTAAGGGGTAATTCTGAAAACAACATTAATAGCAACTGATTATGTAGACTAATTCAAAGATCTACTCTGTATGTTTATTATTTTCAGAGTTATGTTCCAAACTACACTTAAAAATAGACTAATTTTTTAATTTTCCTTAAAAAAACATAATGGTTTACACTCCTCCTTCCCGTAGAACCTCTTAATGTCAAGGAATCTCTAAAAAGGAAATTGAGAATAGAGATTTAAGAACCCACTTAACTTCTTAACCAGAAGCACTAGGCAAATATAAGAACATTAGACAGATTTCC

**>Fragment N (**Natanaelsson et al. 2006; Ding et al. 2012)

**T**TGAACTACAGAAAAATGGGGTAATGATTCCAAACCCTGTGGAAATTGGTCTTAATGAAATAATAAGAGCACATCATGTAGCAGGCTGTTGAACTTTCCGGGGATTTGGTGTTGCTGTCTGACTCACTTTGTCGGTATTGATAGCCATGAGCTAAGAATAGCATTAAGGATTAATGTACCAACAATTAATATACCTTATTTTTAAATTTTAGGCTCCTTCATTGTCATCATCCTCTTGATATTATGCCATGAATATTAAGTGGCACCTTTTCTGGTCTTCAGGAAAGGTCCTGTAGCTGTTCATAAAACTGCTGACTAAAAGCTATGTACTATGCAACCTTCCAAGAATAGAGTGTCAACTAACTGGACATAGGAGAAAGCTGCCTCTACTCCAGGACTCTTCACAAAGTTGATCCAGCTGTAACTAATAATAGTAATAATCACATGTTTTATGGTATCTGACAGAATGTGGCAGTGTCACAGACTACTGACTTGAAGATGATCTTTTTCTATGTTCCTGTATTTCTGAACAGCTAAATTACTTTTCTTCTATATTTTACCTTTTCATAATTTTTCTGAGGAAAAAATACTATTTTAGCTGGCCATTTGCTTGGTAATAAGATAGTTAACAAATTCAAACCTTTAATCATCTGGGTTTTTTTTAATGTGAAAAATTAAATGAGAAAGTTTTTTGCGGTTATGTTTGGTTTGATTTGGTTTCTTACCATTTTATATTTTTCTCTGTTTGAGGCCACAGTGATTACAGTTCTGGTTCCAAAATAAAACTTTAAGTAAAAACTTATACCCTAATAGCCAAGACACAAAGATAATGGATTGCACATAGAGTAAGGAATAAACTTCAGATTTGTGATTTTTGTTTCTAATCCTGATACAGATTTACACTATTTATGAATACATATTTATTGCTTGAAGATATTTGTGAATGGAATGTTGTTATTTTATCTGGGGTTTACCTGCCATTAAATATTAAGGAGTTCTGTAATTTTAAACACTACTCCTTTTACATTTTCTATGTGTAAATAAAACTGCTTAGCATTGTACAGAAACTTTTATTAAAATTGTTTAATGTTTAAAGGGTTTTCCACTGTTTGAGTTTTAAAAAGGACTTTCTGTAAAGACATAGCGTTTGTTCATTTTCAAATCTGATTATATGTATTTTATATATATACATATATATTCACTCGTATAGTGTATAACATAGAAATAAAAATATATAATTGTATAAAGATAAAACAGATTTTAAAATTTTCACATTTA

**>Fragment Q (**Natanaelsson et al. 2006; Ding et al. 2012)

AGAATCTTCTAGTGTTTTATGGGTTTAAGAATTGATATTAGGTGATAATAGATCATTTGTTACTTTGGGTAGCTTTATAACTTAAAAAACTTTGTTTCCTCCTAGCCTGCCAGTATAAATTGGCAGTGGAACGGTATGAATGGAACAAATTGCAAAGTGTGAAGTCAGTAGTACCCATGGTGCATCTTTCTTGGAATATGGCACGAAATATCAAAGTCTCAGATCCCAAGCTTTTTGAAATGATTAAGTAAGTGTTTCCAAAATCACTGTAGTCCCTTTTTTTGTGGGGTGTGGGGATGTGGTGGGAATGTTAACTAATGAACATTTTTCCCCCAAATTATCTTAATTAACTTCAAATGATAAAATGGTCCAGAAGTCTATACAAAAGAGAATTAACATGTTAAAAGTATTTTCTTCCTTTACAAATAAACAATGATTATTATGGTAAAGAGAGGAATAGAGGCATATATACATAGTTCCATGCCAATAAATTCATGATCTGTGCTTTCTCTTGCCCTCTCACCAAATTGATTTTCCAGAACAACTTTAGTTTTAATATTGGAAGTCTGCTGTAGGAACAGAAAGTGATAATTCTATTATAAAACAAATGTTCAGTAAAGAAATTAGCATTTAAAGTGGTGATTTAATTATAATTATTGAATTTTCCAACTATTCTATCTAAATCATCACCTTTTTAGCTTAAGTTCAATTTTAATTTAATAGATTATCAGTTTNNNNNNNNNNNNNNNNNNNNNNNNNNNNNNNNNNNNNNNNNNNNNNNNNNNNNNNNNNNNNNNNNNNNNNNNNNNNNNNNNAGTAACTCATAGTTACTCTCATAGTAACTTCTCATTTTTTAACTTGATTTCCATTCTATTTCTTTCTTTTGGGGTATATTTGTAATAATATGTTTTCACAGACTGCCAAATAACTGTATTTCTCCAAACAGCAGTTTATATGCTTTTCTTTCCCTCTCCCCTTCATTATCACCAATGAAAAGTAACTCTGGTCTGAGAAATGTGGGATGTTACAGATGATACAATTAAGCTGCTGGTTTTTTTCATTGTTACTGTTTTTGAGATATGTGTGTGNNNNNNNNNNCACACAGGAGAGGAAGATTTTTCATGTCTTTGGAATCTGTTTGGCCAGTATCTTTAAAATCACTGCTAGTTTATTACAATTTCAGTTCTATATTGAAAATTTTGTTTAATAAATCTATTTAAACAAAAGCTGAGGCTATAGTAATATGCATATCCACTGTTTTATATAGAGACTCATTCCATTTTTTACATATATCTGAAATTTAAAATAAACATTTGAAAGAGCATATAGTCAAAACTCAGTAATTCTTATTAAAAATGTACACGTTGTCAGTAATTTCTCTATCAAGCTCATTACTTTAAGTATGTTTCTTGAAAATATTCACTGGCTGGTTCAAAATGAACCATTTGTTCTCAGTGCCTTGGATTTTTTTTAGGATTTATGCAGAATGCATCACTTCTGTTTGAGAATTGCGGAGTTTATGTCAGACACTGTTCTTGGCAATGAAGATAGTGTGTAAGACAAAATCTCTTTGAGTTCATATGGTTTAAATTTTAGTGGAGCCAGAGAGGCATATTTTCAGTCAGGTAAATAACTTACCAGTAGTAAGTAAGGTAAGTAAAATAAAATAGGGAAATTTTTTGTTTACAGGATAGATAATACTGCTTTTCCTGAAATGACATCTAAGGGTATGATTTGTGTAGTCATACCTGAATAGTGAAAAGGAGATATTCTTGAGAAGATTTAGGAGTTGAGTAGAACATTTCCTATGAAACATGTGAAAGGCTTAAATTAAGA

**>Fragment R (**Natanaelsson et al. 2006; Ding et al. 2012)

AGTACTAATTTAGAGATTCTTGAGGATATGATGAGAAATAAGAGATGCTTTCTCTTTAGTTAAGAAAATATATGTGATTACTGCAAGAATATATACTAAATTTGATTTCTAACAAAGAATATGTGCTTCTTTAAAGAACCTTGAAATGAATTACTCAGACTTACATGAAATTTGCACATCTAGCTGTGGTAAGGATCCACTGTTCATTCAGTATAGAGCCAGCACAGAAATGGGAGAAAGAGAGTTGCACAGAAACCATCCAGGGAAATTCACCCACTTCTGCTTCCCAACAGTTTGGACAGTGGGGAGCAAGGCCAGGTCGTAATCCACACTCTGTAGAAGGAATCCATGTTTCCAATAGGTTGAAGTTAGTATTACTACCTCCTTTACTTGATCATTTTGATTTCCTATTTTTAGATTTTACAAAAGTGCTTGTTTATTTTCAACTTTAGGAAAATGCTTGGGGAAAAAAAGGAAAATGCTTGGAACTATATAATTTTAACCCTATAGAGTTGGCTATGGGAACAGAAATTGTCTGCATACACTCTGAAAATACTATATTCATTACTCAAATAGGAAAAATAAGCAGTTTCTAAGTACTACACATATTTTTATAACATGCTCTGAAAGAAACTATATTTATAAAACAAACTGTTTCTAGCAAAAATTAATCTTCTCTCAATAAGTACACACCTTAATATCTTGTTATTTTTGTTCCCCCTTCCCTGATCTAAATGTTTTGAATATTATCCTGCCCAAGTATGAGTAAAAGACTTCTGACTTAATCAAAAACTCTTGTTTTAGGCTCTAACAAAGACCTTGCTCCCAAGGATAACTCAAATAGATGACTACACTGAACTCACCTGTGGTTTCATGAGAAATAACAGCTGTCTTGAAGAGCCAGAAATCCTGGATAGTCTCTTTCCAGATCACACCCAGGTGACACTCAGCCAAAGAAAT

**>Fragment DBY4** (MG594247; Hellborg and Ellegren 2003, Fain and Straughan, unpublished)

TGGCAGCCGTGGTGAYAGRACTGGCTTTGGCAAATATGAACGCAGTGGACACAGTCGTTGGTGTGACAAATCAGATGAAGATGATTGGTCAAAACCACTTCCACCAAGTGAACGATTGGAGCAGTAAGTTTTTGAAATGTGTGTTGACTGATGTAAAACTTACTACTTAGTATAACATGTAATAATCATTTGATTTTTAGGGAACTCTTTTCTGGAGGAAACACTGGAATAAACTTTGAGAAATATGATGATATACCAGTAGAGGCAACCG

**>Fragment DBY5** (MG594244; Hellborg and Ellegren 2003, Fain and Straughan, unpublished)

CTTTGAGAAATATGATGATATACCAGTAGAGGCAACTGGCAATAACTGTCCTCCACATATTGAAAGTGTAAGTTGCTTTTGTTTGACGTTTTAACAGTTCTTTGTTACCACTTAAAAACTTCATTAGATTATAAAGAAAGGAGAGTCTGAGACTTTATATTACCATTATGACCAGTGCTGAAACAAAATGTCAAACATAAGGAATCCAAAATTAATCTAATATTAGACCCAGGAGTCTTATTTTCTTAAAAACACGATGATACTTTTAACCTCAAATTATAGCAAATTTGGTTTGTAGTCACAAACCACAAAAGTAGATATTTGGGGGTGGTTTTATTGTCATATTAGTACAAAACAGAAATTGTAACAGCTAAGGCATTTCTAGTTTTATGTGTATTTTACTTAATATTTGATAGTTCAGCGATGTTGAGATGGGAGAAATTATCATGGGGAACATTGAACTTACTCG

**>Fragment DBY7** (MG594246; Hellborg and Ellegren 2003, Fain and Straughan, unpublished)

GGTCCAGGAGAGGCTTTGAAGGCTGTGAAGGTAAAGATTGATTTCCCTCTAAAATAAGTTGTTGGCATAGAAAAAGCTTCATAAATATAAGTTGAGACCTTCCTTTTAAACAATGGCAAATATGTTTTATTTCAGTTCCTTTGAGTGTTTTATTATATCTCATGAAAATGTATTTTTGATGTAATTTTTAATTATACAGGGAATTTTGCTGTAGAGTTGTTTTAAGAGAAAAAAAGAATTCTGGGTCTTTGAGTTTAATTTGTAAATTATACTCATTTCTTAGGAAAATGGAAGATACGGACGCCGCAAACAATATCCTATCTCCCTAGTTCTAGCCCCAACAAGAGAATTGGCTG

**>Fragment DBY8** (MG594245; Hellborg and Ellegren 2003, Fain and Straughan, unpublished)

**C**CCCAACAAGAGAATTGGCTGTACAGATTTATGAGGAAGCCAGAAAAGTAAGTGTGCATTTTACTGATTATTGCCATTCTCATTGATTGTAGGGATGTTTTTATGAAAATCTGATTAAATTATTTGCTTATAGTTTTCATACCGGTCTAGAGTTCGTCCTTGTGTAGTCTATGGTGGTGCTG

**>Fragment SMCY5** (MG594248; Hellborg and Ellegren 2003, Fain and Straughan, unpublished)

**A**AGGATAAGACTTTACGGAAGAAAGGTGAAACCTCATAGGCCAAGAGGCACTGTGTGGGTTAAAGTACAAAAGTGGAAGAGGTGGTTTGTTACTTTGATAGGAGAATTGATTTGTTTTCAGATTTGTTTCTGACTGCCCTAGATAAAGAAGGGGTTTTGTGCCCCCGAGCTGTTCTGATGAAAGAGGAACCAAGAGAGGATGAGAGAGTGATGTCCTCATTGCCCAGAGAGGACTTGAATCAC

**>Fragment SMCY9** (MG594249; Hellborg and Ellegren 2003, Fain and Straughan, unpublished)

**C**AAAGTACGAGTRGCYTTGGAAGTGGAGGATGGCCGGAAGCGCAGTAAGTGATGAGAAATGGAAGGATCCATGGGGAAGCTCTGTTTGTTTTTTTCAGCACTCCAAGCCTCCCTCCAATATTTTTCCTAATATTGTATACTTTTTGCAACTTTACCTCTATCTTCAGGCSTTTGAAGAGCTAAGAGCA

**>Fragment SMCY11** (MG594251; Hellborg and Ellegren 2003, Fain and Straughan, unpublished)

TGCCATGCATCAAATAGAGGATGTCAAGGTGAAGAGGAAATTCTTATTAAAAACAGATACTAGCAAAATCTGAGGTAGCAGGGAAGCTTGAGAACCTAATTGTTCTAAGTAATTTCTTATAATACATCTTTCTTACAATTTATAGAATTGAGTACTATTTGTATCATTGAGTACTATTTGTATCATAATATTCATAGTATATAGCTTCATAACAGTATAGCTTATAGCTTATTATTATTATTCTTATAGCTTATAGCTTGTATACTTAAAGCGGGTATCTTTCCATGCAAAAATAATTTAAAATACTGAACACAGGGCTACATACCTATCTGGCTTTTGGCACTGGAAAATGTGAAGGAGATAGAGTTGTTTTTTGTTGTTGTTGTTGTTTTTTTAATAATTTAGAGTCTTAGGAAGCCTTCGAGAAGAATAAGATGGTGTTAAAGTGGGCCTCAAAGGATAAATTTCAGATAACTAAAGGAAAAGTTGGGTTATTATGGCAGATAAAGGAGTTTTGTTGATATTAAGTAGATCGTGTCTTATTATATGATTGAAGACAAGCGTAAACACCTAAGAGAAAACATAAACAGATTAAGGGCAGGGGTTGTGTAACTAGGTGGCTAAAGAGTCTATCCTTTATCCTATATCTCTGCAGGACGTCCTGGAACAGGTGG

**>Fragment SMCY17** (MG594254; Hellborg and Ellegren 2003, Fain and Straughan, unpublished)

TTTTTGCTCGAGTAGTCGGCAGTATCTGCGGTGAGCCATGGGGTCTACTTGGAGGATACAGTGGAGGTTACAGAGCTGGGCCAGATGTGCTTTTCCTTGCCCCCTTCCTCCAGGTATCGGTACACCTTGGATGAGCTTCCTGCCATGCTGCATAAGCTA

**>Fragment SRY** (AF107021, Meyers-Wallen et al. 1999)

CCCGGTTAGATGTTGACCTTATTTGGGAGTTTTGCTTGAGAATCGGTAGGTTGGGCTTCGGGTGATGGTCAGCGGAGCGGGTTTGAGGCAAGGTGCTGGGCGGAGAAATGAGTATTTTAGAAGCAAAAATCACAGCACCAGAGTCTAGATAATTTTCTGAACGCTTACACTTTCCAACTTCCCTCCGTACGCATCGCCCCCCGTCCCCCCCCCCCCCCCTTTTTTCTGTAAGCGCTGTAATTTTACGCTTCTGCTATGTTCAGGGCATTGAACTGCGATGATCACGGTGCAGCGGTACAACAAAATGCCTTCGGCTTTCCGAGGAAGTCTTCCGACCGTTGGACGGACAATTCAACCTCGAATTATCGGTGTGAAAGCGGAGGAAACGGTAGAGACAGCGGCCGCAATCGCGTCAGACGACCCATGAACGCATTCTTGGTGTGGTCTCGCGATCAAAGGCGCAAGATGGCTCTAGAGAATCCCCAAATGCAAAACTCAGAGATCAGCAAGCAGCTGGGGTACCAGTGGAAAATGCTTACAGAAGCCGAAAAATGGCCATTCTTCGAGGAGGCGCAGAGACTACAGGCCATGCACCGAGAGAAATACCCGGACTATAAATACCGACCTCGTCGGAAGGCCACGGCACAGAAAAGTCACAAATTGCTACCTGCAGCCTCCTCCTCCATGCTATGCAAGCAGGTGCACGTAGATGAGAGGTTCTACCCCTTCACCTACACGGACAGCTGTAGTAGGGCTGCACACACACGAATGGAGGACCAGTTAAGCTGCTCACAACCCATGAGCACAGCCAGGTCGCTGCTGCAACAGGAGTACCACAGCAGCTCCGCAAGCCTCCGTGACAGTCCGGAAACCTTGGCCGCACAGATGTCCGCTGACGCTTCTTTTTACTCTAAGTAACAGCTGGGACTTTCTGACGCTTATTTTCCTTAGTGATTTCTTTACTCTGGCTAACGAAACGTCCTATTCATACTGATTTTGCTATTATTTCATTCAAAATGAGCGTTGGAAACTTGCTTAACATATAAAGAATTACAAAGTATGCAATTGACTCAGGCTGGGTTTATGGCTGCTTAACTTGAGGAGAGAGCACGTTGATACTTTCTGAAACTCCTGGACTTCCAAGACCACTTGTTTTTCTTTTAACCAAACAACTGTTATATCTAATTATAGTTGCTCCATAGACTGAGCTTTAAATAAGTGAAATAATAGGTGCAAGAATATACTAATAATCTGATAATCGTCCTCATAGGTACCACAATTTTCACTCTTTATTTTTAAAATACTTCCTTTATTCATGAGAGACACCCAGAAAGAGGCAGAGACGTAGGCAGCGGGACAAGCAGGCTCCCTGCGGGGGGGAGACCGGTGGGGGACTCGCTGTCAGGAACCCCTCCCCCCCACCCCCCCATGGGGCCCTGAGCCAAAGTCCCACGCTCTACCCCTGAAGCCCCCGGGTGGACCTTCACTCTTTATTTTAAATAAATATTATTTATTATTATATTATTATATTATATTACATTATTTAAATAAATATTATAAATAAATATTTTTATCACA

**>Fragment UBE1Y6** (MG594255; Hellborg and Ellegren 2003, Fain and Straughan, unpublished)

**T**GTGGCAGCATCCAACCTCCGAGCAGAAAACTATAACATTCCCCCTGCAGATCGGCATAAGGTAATGATAGTCTAGAGCTAAGCTTTCACCCTCTCCCAAGGTTGAGTCTCCCAAATTTGTCCTCTTTTATAGATCCTGAACCTCTTTATAGACTGCATATGCTGGTCATCTTCTGTCCCTGCAGAGCAAGCTGATTGCAGGAAAGATCATCCCTGCCATTGCCACAACCACAGCAGCTATAGTTGGCCTCGTGTGTCTGGAGCTATACAAAGTGGTGCAGGGACACCAGCAGCTTGAGTCTTACAAGAACAGTTTCATCAACTTGGCC

**>Fragment UTY11** (Hellborg and Ellegren 2003, Fain and Straughan, unpublished)

**A**AAATTGTATTTGGTACACTGAACATTTAAGAGTCTTTAAATCTGCCAATAGTGGCATCATCTAAACATGTGCTAAATAGGCAGAGAGAGACAGAACTAAAAAAACTAAAAAGGGAAAAAAGTTTCTGCTAGTGTTTTAAAATATTAAAAATAGTAAAAGGTATTTCAAAGTTTATTTTTTCAATCTTACCTAACTTCAGAGCTCCAGCAAGGCCACGTATTACTGTAACTGGGTTTTTTGGATT

**>Fragment ZFY Intron** (AB622147; Tsubouchi et al. 2012)

GGCATCTCTTTACTATGCTTGGTTTTCACATGTGTTTTTAAGTTAGAAGAGTCTGCAGACCTATATTCGCAGTATTGGCACTGGTATGGCTTCTCGCCAGTGTGGATTCGCATGTGCTTTTTGAGCTCTGACGGGTGACGAAAACCTTTACCACACTCCACACAAATATGAGGAAAGTTCTTGCTGTGAACCGCCAAAAGGTGGCGATTCAATAAGCCTTGTTCAGCTGTCTCATATTCACAGAATTTACACTTGTGCATTTTGTTGGTTCCCTTTTCCTTATGCACCATTTTGTGTGTAAACAAAGCCCCAGCGTGAGAGAAATGCTTCCCACACTCCTCGCATTCGATAGACTTTTCTGCCTTGCTTGTTAGCTTGTGGCTCTCCAGGTGGTTGTGTAAACTTATCTTCTTATTTGTAGTGTAATCACAGTCAGTACAGCGGTACTTCTTCTTGGAAAGGTGTTCAGGATGGTTTTTCATGTGCCTTTTCAAAAAACCTCTCGACTTAAATTTTTTTCCACAAATCATGCAAGGATAGACAGTCAAGGGATGTCCATCAGGGCCAATAATTATTGCTAAGAAAGGGAAAGAAAAGAACATGAGTGATCAAACCAAGTTCTGCTTTGGTTCTTCAACCACTTAAAGTGTGTGCTCTGAAAATAGTGCTAAAGTACTCCTAAAAGCTAATATTCTTTCGTTTTTCAACAAAAGAGGATAGAAAAACCCAGTAATGAAGAAAATGTTGTTCAGAAACTTCATTCAATGTAACTACAAATCTCATTTCCAGTTTAAAAGACATTTAAAGCCATATATAGGGGTGTCTGGGTGGCTCAGTGGTTGATTTAGCAGTGCTAAATCTGCTTAGTTCTTAGTATCTGCCTTAGTTCAGGTCCTGATTCTGGGATTCTGGGATGGAGTCCTGCATGTGGCTCACTGCTGAGAGCCTGCTTCTCCCTTTGCCTACGTCTCTGCCTCTCTGTGTCTCTTATGAATAAGTAACAATAAAAAAGCCATATATACTTGGGAATCTATAGAAAATAATTCATAAAAACAATTCATAAAACCTTTCAGTTTCATTACTGCAAACCAAA

**>Fragment MS34A** (AF032441, Olivier et al. 1999, Sunqvist et al. 2001)

GATCCCTGGGAGGATTACTTTTCAACTCAGATTTAAGCATTCTCTTCCAGTTGGTCCAAGGCATGCTCAAAGAAATAGGTGCAACCTGTAGATGTGGCCTCTTCCCTTCTGGGAGATGGATAATATGCCACAGCCTGTTCGCTCTCGGCAGGCACTCCAATTTACCATGGGGATAATCTTGGATTGCCTCCCTAATTCCTTGGGATGGATAGTCGGTGCCAAGGAGAGTGTTACTAGCCATTCCTGGCCGAGTGGCCTCTCCTGTACCCCAGCTCTCCCATATCCATGTANNNNNNNNNNNNNNNNNNNNCACACATGCATCCATGTGTTCATATACACAATGATGGTTCATACAAAACTTTGGAATATCTACCCCACATCATCAGACACTATGGCAAAAGGGGACCTAGAGCATCCACTCGTCAACCTGATGACCTACCCTCAATGGAGCAAAGAATTCAATCAGAAGAACAAGAAAGTGTAGACAGCCACCCAACATCAAGTTCTTTGTCTTGTATTCACGAAAACTAACTTTACACTTAAGTAAAATGCTCAGCAATTTCTACATTTCTCGGACCTCTCAGCAGCCAGAGTGTCTACCAAAATATAAGTGCAAGAGGCTACATGGGATTACAGACAGCTAGTGTATGAAGGGATAATGTCAGGTGCTCATCTTGCCTTTCCTACTTGCTCTTTTGTGATACCTAGAACAATGATATGAAGGGTGGAGCTACAGCCACCTTTCTGGACCAAGAAGATTGGGAAGACTAGAACCATGGAACCATTGCCAACCCCCAGGTATTTCTGGACTTTTACTTCATGTTGAAAGTGAATACACTTCAGTCATTGTACACACTTACATCTGTGTGCTTCATGTTCTGTTATTAAGAGATGGATGTAGCTCATAACTATTACAGAAGATC

**>Fragment MS34B** (AF032441, Olivier et al. 1999, Sunqvist et al. 2001)

GATCCCTGGGAGGATTACTTTTCAACTCAGATTTAAGCATTCTCTTCCAGTTGGTCCAAGGCATGCTCAAAGAAACAGGTGCAACCTGTAGATGTGGCCTCTTCCCTTCTGGGGGATGGATAATATGCCACAGCCTGTTCGCTCTCGGCAGGCACTCCAATTTACCATGGGGATAATCTTGGATTGCCTCCCTAATTCCTTGGGATGGATAGTCGGTGCCAAGGAGAGTGTTACTAGCCATTCCTGGCCGAGTCCTCTCCTGTACCCCAGCTCTCCCATATCCATGTANNNNNNNNNNNNNNNNNNNNNNNNNNNNNNTGCATCCATGTGTTCATATTCACAATGATGGTTCATACAAAACTTTGGAATATCTACCCCACATCATCAGACACTATGGCAAAAGGGGACCTAGAGCATCCACTCGTCAACCTGATGACCTACCCTCAATGGAGCAAAGAATTCAATCAGAAGAACAAGAAAGTGTAGACAGCCACCCAACATCAAGTTCTTTGTCTTGTATTCACGAAAACTAACTTTACACTTAAGTAAAATGCTCAGCAATTTCTACATTTCTCGGACCTCTCAGCAGCCAGAGTGTCTACCAAAATATAAGTGCAAGAGGCTACATGGGATTACAGACAGCTAGTGTATGAAGGGATAATGTCAGGTGCTCATCTTGCCTTTCCTACTTGCTCTTTTGTGATACCTAGAACAATGATATGAAGGGTGGAGCTACAGCCACCTTTCTGGACCAAGAAGATTGGGAAGACTAGAACCATGGAACCATTGCCAACCCCCAGGTATTTCTGGACTTTTACTTCATGTTGAAAGTGAATACACTTCAGTCATTGTACACACTTACATCTGTGTGCTTCATGTTCTGTTATTAAGAGATGGATGTAGCTCATAACTATTACAGAAGATC

**>Fragment MS41A** (AF192268, Olivier et al. 1999, Sunqvist et al. 2001)

GTGGATCCTTTTTTACCTGGTGCACCTGATAATTGTCGAGGGTGTTAAAATAATCGAAGCCACAGACTGTCTCCTCCAATATCTCCTGGATGCGGCTCTACAAACACAGTGCCCTGTTTATAATCAAGAGACAATACGGTGCTCTCTGGACATCACCCAAGGGGACAGCCCCACTAGAAACTGTGTTCTCTAGTTAATGTCACAGGGGCAGCTGAAGAAACATCTGGCAAGGAGGAAGCCAGATGAAAGAGACTCCTGGGCTTATGATTAACTTATTGGAAGTTGAGCTTGGTCCCTAGCACTCACGTTCTCATCTTGCCTGTCATGACCTGGGGTTTGAACACAAAGATGTGTCAATCTTCCAACATCTGGCAGCTGGTTTCTGCCCTGGAAGTGTATATTCCTCTAATTTTCCCCTCTANNNNNNNNNNNNNNNNNNNNCACATACACACACAGATACACACACAGATGGGTAAAGGGTCTCACTCCACTCAGCTGTGATGGTAGTGTGGTCTGCTCACCAGTGAGCTGCCCCAAGTTGCCTTGCTGTACCTGTTGATGTCTCCTGACACATAACAAGAAGCTTCAGAGAAGAGGGTCGAGCAGGCGTACAATGACATGAAAATCCTCATTCTGGACTTTTCTGCACCCAGAGCACCTGAAAGAAGGAAACCAGCAACAGAGCAACCTGTTCTCTTAACCATGTTGAGTTAAATGAGCTGGAAAGCAGGAGGTTCTAGAATGCAGGGTCCTGGTTACCTGAGTTCTAAAGCAGTTGTGGATGTCACCAAGGAAGTGACGTCACTTTTTCAAGATTCCAGCATCTGGGGCTCTTCTTTCAGTCAGACCTATCCAGAACCCCTTCTTGCAGATGACTGCTTACCCCTGTCGTCTCCTTAACTGCCTCATCAGCCAACAAGGCCATTGCCATAACTCTCTGTTAATTCCAGTAAGGTCCTAACTTTTAGGAGACAGAGCTGAATCCAGACCTTTAGTTTTCTGCTTGGCTTCTCAGTTCTGTGTTGTATCTAACTCACGATGTCTGTCATATAGGTCCCTGTTTCCCAATTAGCACAGTGGTAGAATCAGA

**>Fragment MS41B** (AF192268, Olivier et al. 1999, Sunqvist et al. 2001)

GTGGATCCTTTTTTACCTGGTGCACCTGATAATTGTCGAGGGTGTTAAAATAATCGAAGCCACAGACTGTCTCCTCCAATATCTCCTGGATGCGGCTCTACAAACACAGTGCCCTGTTTATAATCAAGAGACAATACGGTGCTCTCTGGACATCACCCAAGGGGACAGCCCCACTAGAAACTGTGTTCTCTAGTTAATGTCACAGGGGCAGCTGAAGAAACATCTGGCAAGGAGGAAGCCAGATGAAAGAGACTCCTGGGCTTATGATTAACTTATTGGAAGTTGAGCTTGGTCCCTAGCACTCACGTTCTCATCTTGCCTGTCATGACCTGGGGTTTGAACACAAAGATGTGTCAATCTTCCAACATCTGGCAGCTGGTTTCTGCCCTGGAAGTGTATATTCCTCTAATTTTCCCCTCTCNNNNNNNNNNNNNNNNNNNNNNNNNNNNNNNNNNNNNNNNNNNNNNNNNNGATACACACACAATGATGGGTAAAGGGTCTCACTCCACTCAGCTGTGATGGTAGTGTGGTCTGCTCACCAGTGAGCTGCCCCAAGTTGCCTTGCTGTACCTGTTGATGTCTCCTGACACATAACAAGAAGCTTCAGAGAAGAGGGTCGAGCAGGCGTACAATGACATGAAAATCCTCATTCTGGACTTTTCTGCACCCAGAGCACCTGAAAGAAGGAAACCAGCAACAGAGCAACCTGTTCTCTTAACCATGTTGAGTTAAATGAGCTGGAAAGCAGGAGGTTCTAGAATGCAGGGTCCTGGTTACCTGAGTTCTAAAGCAGTTGTGGATGTCACCAAGGAAGTGACGTCACTTTTTCAAGATTCCAGCATCTGGGGCTCTTCTTTCAGTCAGACCTATCCAGAACCCCTTCTTGCAGATGACTGCTTACCCCTGTCGTCTCCTTAACTGCCTCATCAGCCAACAAGGCCATTGCCATAACTCTCTGTTAATTCCAGTAAGGTCCTAACTTTTAGGAGACAGAGCTGAATCCAGACCTTTAGTTTTCTGCTTGGCTTCTCAGTTCTGTGTTGTATCTAACTCACGATGTCTGTCATATAGGTCCCTGTTTCCCAATTAGCACAGTGGTAGAATCAGA

**>Fragment 35.4** (AY466399, Bannasch et al. 2005)

TCTACCTTTCCTCCATCCGCAGAACAGGTATTACTTAGAACTCCAGGCAGTAATACAAATGAAGCACCTGCCGTGTGTGTGTGTGTGTGTGTGTGTCTGTCTGTCTGCTCAATTTCTCTATCACCTAAAATAGCGGCCCTTCAAATATATCCAAAGACTAGGTCTCCAGGTGTGGAAGGACAGAGCTACACAGAACCTGGCAATGGATTAAACTGGGGAACTGAGGAAGAAGGAAGAATCAAGGATGTATCCAAATGCCTGACAAAGGTAATCAAGCAGAAGAATATCAAGATGAGTCTGT

**>Fragment 79.2** (AY466397, Bannasch et al. 2005)

TTTTTGGGTTTTTATAAAATACATCAACAGCCTTATCTACCTTCCCTGCCAAGATATTGGACAGATGTATGTCAAATGAGGATAACTCTCAGGAGCATGGTTTATTCTAGTCTTTACTTTTCCACTCAGACTTAAGCATTCTATTCCATTTGGTCCAGGGCAGGGGAAAGGAAACAGGTGCAACCTGCAGATATGGCCTCTTCTCTCCAGGGAACTGGAGAATATGCAACAGCCTGTTGGCTCTTGGGGAGCCCTCCGATTCACCATGAATATAATCTGGGAATGCCTCCCCAATTCCTTGGGGTGGAGTGCCAGTCCCTATGGAAAGTGGTGTTATCCAGTCTTGGTCATCAGGTGACGCCCCTGCACTCAGCTCTCCCTTGTCCATGNNNNNNNNNNNNNNNNNNNNNNNNNNNNNNCACATCCATGTGTGCACACAATGATGGCTCACACAAAACTTAGAAATATATACCCCACATTATATGACACTCTGAGACCTGCCCTCAACT

**>Fragment 79.3** (Y466398, Bannasch et al. 2005**)**

CTCCCTGGACATCACCCCAAGAGGATGCCACATTAGAAACCATGTTCTCCAGTTCAGCCCAGAGGGGCACCTTAAGAAATATTTGACAAGGAGGAAGCCATATGAAAGAGACTCTTGGGTTCATGACTAACATAGTCAGGGTTGAGCTTGTTCCCTAGCACACGCCTTCTCATCTTGCCTGCTATGACCTGGAGTCTGAAAACAAAAGATGTCCAATACCTGGCAGTTAGTTTCTGCCCAGGAAGGACATTTTCCTCTCCTCTGTACCCTGTAGACAAGCAAAGACACAAANNNNNNNNNNNNNNNNNNNNNNNNNNNNNNCACAGTGAGGGGTAAAAAGTTTCAAACCGCTCAGCTGAGATCCAATGACCAACCAAAGTTTTTTTTTCCAACCAAAGTTTTACTTGATGTTTTACAAAATTTTTAAGGATATTTATTAATTTCTTGAAGATAGAGACTAAGCAGGGTGAGCAGAAGAGGGAGCGAGAGAATCAGGGAGCCCAGTATAGGGTTCTAACCCTGGATTCTGGATCAGTGCTGTAATGAGGTGAGAATGGAAAGAGGGATCTT

**>Fragment 650** (AF005413, Olivier and Lust 1998)

GTCCTGGGTTGAAGCCCTACATTGGGATCTCTGCTCAGTGGGGAGCTGCTTCTCCTGCTCCTTTTGCTGCTCACTCTACTGATTCTCTCATTCTCTCTCCAAGAAATTAATAAATACCCTTAACAAGTTTTGGTAAAATATCAAGGAAAATTCTGATTGCTGTTTGGATCCAGAAGCAAAACCTTGATACCTCTACAGAGCACTTAAGGAGACAAAAGGGATTTTGGCTGAGGAGTGACTCTCAGAGTCCAGAATCTCATAGATCTCTTGATTTCCATTGAGAGGGCTCATTCGAGGACTTCCTGTGATGAGGCAACTGAGTCAAATGGAGCACCAGATGGAGTCCCTAACCCGAAACTGGTCCCTAACCCTAACCTGAACACAAACCCCAACTTGATCCCTAATACCTAATCTAACATGAACCTTGACCCTAACCCTAATATGAGCCATAATACAAACCAGACCCAAACTCAAAACCCAACCCTAATCCAAACTCTAACCTGAACTTTAGTCAAACCCTAACCTAAAACCTAATCTGACCCAAACCTGAAAAAGAAACCATAAACCTAAACCTAACCCTAAACCCTAATCCTTAACCCTAACTAGAACAAGTGCCCTATTCTGAACCCCAAGTCTAACACTAACCCGAACCCAGGAC

**>Fragment 990** (AF005414, Olivier and Lust 1998)

GTCCTGGGTTCTCCATTGTGTTCCACTGATCTGTGTGGCAACTTTTGTGCCAGTACCATATCCTTTTGATGATGACTACAGCTTTATAATATAGTTTGAAAACTGGAATTGTGATGCCTCCAGGATTGGTTTCCTTTTTTAACACTCCTCCGGCTATGTCAGGGTGTTTTCTGGTTCCATGCAAATTTTAGGGTTGTTTGTTCCAGGTCTGTGAAAAATGTCCATGGGATTGCATTGAATGTGTAGATTGCTTTCGGTAGTACAGACATTTTTACAATATTTGTCCTTCAGATCCATGAACATGGAATCTTTTTCCATTTCCTTGTGTCATCTTCAATTTCTTTCATTAGTGCTTTATAGTTTTCAGAGTACAGATCGTCTACCTCTTTGGTTAGGTTTGTCCCTAGGTGTCTTATGGGTTTTGGTGCAATTGTAAGTGGAACTGATTTCTTGGTTTCTCTCTCTGCTGCCTCATTATTGGGGTATAGAAATGCAACAGACTCTATACCTTGATTTTGTTATCCCGCAACTTTACTGAACTCAGGTATTAATTGTGGGAATTGTTTGGAGGAGTCTTTTAGGTTTTCTACATAGAGTATCATGCCACCTGTGAATAGTGGAGGTTTGACTTCTTCCTTGATGATTTGGAAGTCTTTTATTTCTTTGTGTTTTGTTGTTGTTGTGACTATGACTTCTGGTCCTACATTAAATAACTGTGGTGAGAGTGGACATTCCTGTCTTATTCCTGACCATAGAGGAAAAGCTCTTGGTTTCTCCCCACCGAGGATGACATTAGCTGTGAGTCTTTCCTAAACGGCCTTTATGATGATGCGGTATGTTCCCTCTATCCCCACTTTCTAGAGGGTTTTTTTTATCAAGAATAGATGCATCTACTGAGGGAACCATATAGTCCTTATCCTATCTTTTATTAAAGTGGCGTATCATGTTGATCGATTTTGAAATATTGAACCAACCCTGCAACCCAGGAC

**>Fragment AMELY Exon** (KC763835, Yan et al. 2013)

GGCACCCTGGTTATATCAACTTCAGCTATGAGGTAATTTTTCTCTTTACAAATTTTTTTTCTTTTTTTTTTTTCTTTACAAATTTTGACCATTGTTTGAGTTAACAATGCCCTGGGCTCTGCAAGAATAGTGTATTGATTCTTTAAGACATTTCTTAGTCCCACCTTTTCAGTTTATATTACCAGATTTCTCCTTTGATGAGTTGCCTCAAGCCTGCATTGCCTCTGCACACTCTTACTTGGCCTCTCTGACTCTGTGTCTCCTTCTTAAATGGCTATGAAGTTATTTATCATAAACTACTGCTCAGGGTGTGGCTGCATAGTAGGACAGAAAGTAAACTCTGGCTGAAGAGCTTTGTTCTATTCTAGTCCTATAAAAGATGGGGGAGTCAAGTCAAGTCAAGTTATTTAGGTTCCTTTCCAGCTAGAAACTCATGATTCTAAGATTTTCACAGTCTGTGTCTCCCTTCTTTGTCTTTACTGAAAGATCAATGATCAATAAGTCTATGAGACATTCTCACAGAAATGGAAAGCATGAGGCTCTCCTTAGCAATATACTAACCACAGGAAGTTGGGCAAATCATTTTTGTCTCAGGAATTTTGGTTTCCTCATCTGAAGAAAGGAAATTGTTTTAAATGCCCACACTTCAAATAGTTTGAGGAGCAAAATAGTTAAGTAATATAAAAAGTTCTTCATCAGATATAACTTTAGCAAAGTTATAATTGTTTCTTGTACTGATATAAAATGCTATACTACTACTTGAAGGCATCTTTCATCTCGTTAACCTTTTCACTATGACATAAAGACAAAAAGGTGTATAATTAATTTTATATCACATTAATTTTTTGTTTTGTTTTGGTCTTGCAAAGAAGAGCAGGAGCTATGGAATTAAACAAAAACACCCCAGTAAGACTATGAACTTCCTTAATTCCTCATGACTTTGTCTGTAAAAATGACAGTGTTGATATTTGTCTCAGCATATATGAGAGAGCATGAAATGTGTATGTGATTGACATAATACACTGCCTAACACATAAGAAGTACCTGGGAAATGTTTACTTTATTCTTTCTTCTGTAGAACTCAATTTTCAGACTATCAATATTGATGAGACTTTAATAGTTTGTATTTTATGCAGCTAAATTAAGGCAAATGAATTCTAACATCTCTTTCTCTTAAGGTGCTTACACCTCTGAAATGGTACCAGAATATAAGGCATCCGGTATGCAGACTTTTTGTCCTTTATTCCCTTAGAATATAAAATGTATTTCAATTTTCTTTTAAGTGAAATAATGTACCTATGCCACATACAGATCCTAATGGGAAATCTAGTTTGTAAAATGTCATATCTATGTGCACAGTTAGAAATTTCTCTACAAGGGAAAGTGGATAAATATTCATACTATCATAATGACAAAGAAAACATGACTATATCTCCTGTTGGAAGTCAGTTTGACTGAGTCACTGTAACTGAGTCAGTTGTGTGAGCTAATGATGAACCTGATTCTTTGTTTCCCACCAGTACCCTTCCTATGGTTACGAACCCATGGGTGGTTGGCTGCACCACCAAATCATTCCCATGCTGTCCCAGCAAAATCCCTCGAATCAAGCCCTGCAGCCTCATCATCACATCTCCATGGTGCCAGCTCAGCAGCCTGTGGTCCCCCAGCAACCAATGATGCCAGTTCCTGGCCAACACTCTATGACTCCAACTCAACATCACCAGCCAAACCTCCCTCTGCCTGCACAGCAGCCCTTCCAGCCACAGCCCGTCCAGCCACAGCCTCACCAGCCCATTCAGCCACAGCCACCTATGCACCCCATCCAGCCCCTGCTACCACAGCCACCTCTACCTCCGATGTTCCCCATACAGCCCCTTCCCCCCATGCTTCCTGACCTGCCACTGGAAGCTTGGCCAGCAACAGACAAGACCAAGCAGGAAGAAGTGG

**>FRAGMENT ZFY Exon** (JX475923, Gilbert and Silversides, unpublished)
ATGGTGGCTCCTCCGGAATGACCATCGACACGGAGTCCGAAATTGATCCTTGTAAGGTGGATGGCACTTGCCCTGAAGTCATCAAGGTATACATTTTTAAAGCTGACCCTGGAGAGGATGACTTAGGTAAGAAGAAAGCTTCAGCATATTATACATCCTGATCAGATACACTTCAATTTGATTTCTTTTGGGTGGTTTAGATTGAGGACGTTTGATACTTTTTCTCTTCTCCTCTTTTCTCTTTCTTGCTCTTTTTAGATCGTATTTATTTGTTTGACAGAGAAAACACAAGCAGGGAGAGCAGGAGAGGGAGAAACAGGCTCTCTGTTGAGCAGGAAGCCTGATGTGGGGCTCTGTGATTATGGGATCATGAACCAGGCCAAAAGCAGACTCACCGAAGTGACTGAGCTACCCAGGTGCCCCTGAAATCTTTTCTGAAGAGAACTTTTAAGCAAGAATTTTAGGGCAGCCCCGGTGGCTCAGCCCGGTTAGTGCCGCCTTCAGCCTGGGACATGATCCTGGAGACCCGGGATCGAGTCCCACATCGGGCTCCCTGCATGGAGCCTGCTTCTCCCTCTGCCTGTGTCTCTGCCTCCCTCTCTCTTTCTGTATCTCTCATTAACAAAAAATAATAATAATAATAATCAAGAATTTTATTGTGGCAGTAGAAAAATATATAGAAAGTGTAATAAGTTAAACAGGTGGAAATGAAGTTTTCACATTAAACTTGGAGCTGTCATCTTCCCTGATAGGTGGCACTGTGGACATTGTGGAGAGTGAGCCTGAGAATGACCATGGAGTTGAATTACTTGATCAGAATAGCAGTATTCGAGTTCCAAGGGAAAAGATGGTTTATATGACTGTTAACGATTCTCAGCAAGAAGATGAAGATTTAAGTAAGTAGGTGCCTTTGTTGTGGGAGAAAATTTTGTTTCTGTTTACAGTAATATATGAAAATAATGTTAATGAGTCTGTAACATTTGAAACTTGTTTGTCTGAATTCAAACAAGTATCTTTTTCTGTCTGACCTTTGTAAACCTGATTATAAACGTTTTACTAATTTTCTAATTAGATGTTGCTGAAATTGCTGATGAAGTTTATATGGAAGTCATAGTAGGAGAGGAGGATGCAGCAGTTGCAGCAGCAGCAGCAGCTGCTGTTCATGAACAGCAGATGGATGACAATGAAATAAAAACTTTCATGCCAATAGCATGGGCAGCAGCTTATGGTGAGTCACATACAGTAGCTCAAAGGATTGCATAGTTGTGGAATACGAATTCACCATTGAGAATAATTTCAGTAGTTTCAGATTTCAGAAATAGGAAGTATCGATATAGTGATTCAAGGACACAGAGAATGTATATTCTGCCTATTTTTGCCTTTCATTTTATTATTATTTTCAAGATTTTGTTTGTTTATGAGAGACACAGAAAGAAAGGCAGAGACATAGGCAGAGGGAGGAGCAGGCTCCCTGTGAGGAGTCTGATGTGGGACTTGATCCCAGGATCCCAGGATCACACCCTGAACCAAAGGCAGAAGTTCAACCAATGAGCCACCCAGGCATCTCTCCCTTTAATTTTAATAGGAATTTACTTCGCCTATGTTATCTAAGAGATAGACATTAAAGGTGGAATTTTTATCTTCAGTATCTGGGTCATGATTTTATAATTTTGAGTGTCATCAAGGTGAAAAATGCTGCTACATTTGGAAGCTAGGCTTTTCATGTTGTTCAGTACATAATATGCCCTTATAGCGCATTACGTGGATAGGGAGTTTGTCATTCATGAGTATCGTGGCTTTTCATTGTTATGGTTAATAAAGAATCCCTAATTCTTTATAAAGAATACTGTATAATATTGTTTTTTAATACACATTGTTAGGTAATAATTCTGATGGCATTGAAAACCGGAATGGCACTGCAAGTGCCCTCTTGCAC
